# A model-based hypothesis framework to define and estimate the diel niche via the ‘Diel.Niche’ R package

**DOI:** 10.1101/2023.06.21.545898

**Authors:** Brian D. Gerber, Kadambari Devarajan, Zach J. Farris, Mason Fidino

## Abstract

1. How animals use the diel period (24-hour light-dark cycle) is of fundamental importance to understand their niche. While ecological and evolutionary literature abound with discussion of diel phenotypes (e.g., diurnal, nocturnal, crepuscular, cathemeral), they lack clear and explicit quantitative definitions. As such, inference can be confounded when evaluating hypotheses of animal diel niche switching or plasticity across studies because researchers may be operating under different definitions of diel phenotypes.
2. We propose quantitative definitions of diel phenotypes using four alternative hypotheses sets (Maximizing, Traditional, General, and Selection) aimed at achieving different objectives. Each hypothesis set is composed of mutually exclusive hypotheses defined based on the activity probabilities in the three fundamental periods of light availability (twilight, daytime, and nighttime).
3. We develop a Bayesian modeling framework that compares diel phenotype hypotheses using Bayes factors and estimates model parameters using a multinomial model with linear inequality constraints. Model comparison, parameter estimation, and visualizing results can be done in the Diel.Niche R package. A simplified R Shiny web application is also available.
4. We provide extensive simulation results to guide researchers on the power to discriminate among hypotheses for a range of sample sizes (10 to 1280). We also work through several examples of using data to make inferences on diel activity, and include online vignettes on how to use the Diel.Niche package. We demonstrate how our modeling framework complements analyses that are commonly used to investigate diel activity, such as circular kernel density estimators.
5. Our aim is to encourage standardization of the language of diel activity and bridge conceptual frameworks and hypotheses in diel research with data and models. Lastly, we hope more research focuses on the ecological and conservation importance of understanding how animals use diel time.

## Introduction

The niche is an integral concept in the study of animal ecology and evolution. As the multidimensional characterization of the resources a species needs to survive and reproduce (Hutchinson, 1957), “time” is a fundamental resource needed by species to carryout their life history strategy (Pianka, 1973; Kronfeld-Schor and Dayan, 2003). Yet, research on how animals use time within the diel period (24-hour light-dark cycle) has been a neglected focus of study (Anderson and Wiens, 2017; Gaston, 2019; Kronfeld-Schor and Dayan, 2003).

Among mammals, diel activity has been viewed as a fixed behavioral trait (Levy et al., 2019) constrained by evolution, morphology, and physiology (Hall et al., 2012; Anderson and Wiens, 2017). Major shifts in a species’ diel niche (e.g., nocturnal → diurnal) have been predicted from ecological theory (Schoener, 1974a,b) and observed (Kronfeld-Schor and Dayan, 2003) to be rare. Partly for these reasons mammals are often attributed a single diel niche (e.g., diurnal; Mittermeier and Wilson 2009; Nowak and Walker 1999). Such a categorization makes the implicit assumption that all individuals—across populations and regardless of environmental context—would maintain this diel niche to optimize fitness (e.g., via foraging, predation risk, and reproduction) based on their morphological and physiological traits.

However, recent studies have shown flexibility in mammals’ circadian regulation of their physiology (Riede et al., 2017; van der Veen et al., 2017) which allows for more adaptability in diel activity than previously thought (Rivera et al., 2022; Gaston, 2019). Further, there is growing empirical evidence that many mammal species alter their diel activity in response to environmental context (abiotic and biotic), including temperature (Gallo et al., 2022; Hut et al., 2012), season (Hut et al., 2012; Farris et al., 2015), anthropogenic activity (Gaynor et al., 2018; Moll et al., 2018), landscape feature (Gallo et al., 2022; Rivera et al., 2022), and intra- and inter-specific competition (Cunningham et al., 2019). Translating whether these changes in activity can be characterized as a shift in an animal’s diel niche (e.g., nocturnal → diurnal) depends on how we define the possible phenotypes.

The traditional diel phenotypes for tetrapods are diurnal, nocturnal, and crepuscular (Anderson and Wiens, 2017). A species is commonly described as such based on the majority of of their activity occurring in a single distinct period of light availability (daytime, nighttime, or dawn/dusk, respectively). Less often considered, but an important complementing diel phenotype, is cathemerality (“through the day”; Cox et al. 2023a,b), which is an even amount of activity across the entire diel period, or a large amount of activity across multiple time periods (e.g., day and night; Tattersall 2008). For many studies, there is little consideration for what constitutes a species being assigned one of these diel phenotypes.

Commonly, researchers plot activity curves and make decisions based on their visual interpretation (e.g., Ridout and Linkie 2009). Studies that consider how species change their probability of activity through the day (Gallo et al., 2022; Gaynor et al., 2018) often do not assess switching among theoretically motivated diel phenotypes (Hut et al., 2012). We see several issues with these approaches. Foremost, without explicit definitions, we cannot propose hypotheses about diel phenotypes that are directly evaluated with empirical observations. Second, the lack of explicit quantitative definitions of diel phenotypes means that comparisons across studies may be inappropriate, because researchers may be operating under different definitions. This clearly complicates the evaluation of diel niche switching across studies. Third, qualitative interpretations of diel phenotypes are without measures of uncertainty describing how confident we are that a species is using a given diel phenotype.

Research on the diel activity of wild animals is becoming increasingly prevalent (Figure S1). This is partly due to the growing use of trail cameras (Gilbert et al., 2022), which can continuously sample the spatio-temporal activity of small to large non-volant animals throughout the 24-hr period (Frey et al., 2017). Recently, several thoughtful conceptual frameworks have proposed multiple lines of inquiry focused on diel niche hypotheses (Gaston, 2019; Gilbert et al., 2022). To help move animal diel research in the direction of advancing our understanding related to these conceptual frameworks, we need to connect diel terminology in ecology and evolution with hypotheses that can be tested using empirical data.

We propose a quantitative framework for defining diel phenotypes and probabilistic models that can be evaluated with empirical data and provide a measure of model/hypothesis uncertainty. Importantly, we propose a framework that is appropriate even at relatively small sample sizes. This is critical for being able to evaluate the diel niche of rare and difficult to detect species. Further, we offer multiple hypotheses sets that accomplish alternative objectives.

We implement our framework in the new Diel.Niche R package (R Core Team, 2023) available on Github at (https://github.com/diel-project/Diel-Niche-Modeling). The fundamental elements of the package are 1) defining diel phenotypes, 2) model comparison among diel hypotheses using Bayes factors, and 3) estimating probabilities of twilight (dawn and dusk), daytime (daylight hours), and nighttime (night hours) activity. Users may use all three elements, or choose to estimate probabilities outside of this framework and only use the package to define their results according to preset diel phenotypes. Further, users can modify existing hypotheses and also define completely new hypotheses. For non-R users, a visual online Shiny application is available for simple analyses (https://shiny.celsrs.uri.edu/bgerber/DielNiche/). We hope this work will encourage consistency in definitions for appropriate comparisons across studies.

## Explanation of the method

### Defining diel phenotypes

We define diel phenotypes based on the activity probabilities of the three fundamental light availability periods: twilight (tw), daytime (d), and nighttime (n). We refer to these time periods as ‘diel periods’ throughout. The sample data is thus a vector of observed independent detection frequencies in each period, as **y** = [*y*_tw_ *y*_d_ *y*_n_], respectively. These frequencies depend on the respective probabilities of *p*_tw_, *p*_d_, and *p*_n_.

We translate each diel phenotype as a probabilistic multinomial model with linear inequality constraints on the probabilities (Heck et al., 2019) following the implied relationship under the given phenotype. Specifically, we define inequality constraints by matrix **A** and vector **b**, such that **A*θ*** *≤* **b**, where ***θ*** = [*p*_tw_ *p*_d_]. The sum of the three probabilities is always one, such that there are two free model parameters, where *p*_n_ is derived as 1 *− p*_tw_ *− p*_d_.

As an example of how constraints are used within this modeling framework, let’s define a phenotype where we are interested in whether an animal is primarily diurnal, such that the majority of its activity occurs during the daytime. This phenotype is general without any specific constraints on how much activity occurs during the twilight or nighttime. Simply, the probability of activity in the daytime is greater than the activity probability during twilight or nighttime. We can define this phenotype as a set of two inequalities,

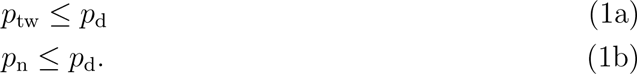

To translate these into the above inequality setup, we redefine the Eq. 1a as,

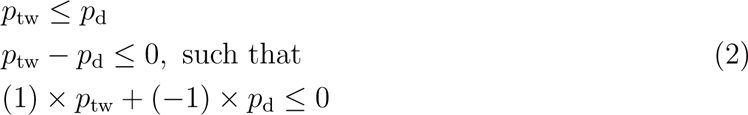

and, given that *p*_n_ = 1 *− p*_tw_ *− p*_d_ according to our earlier definition, the Eq. 1b is,

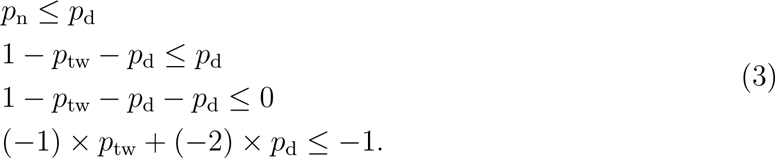

The constants within the parentheses on the left of the inequality sign are packaged into matrix **A** and the constants on the right are packaged into vector **b**, such that

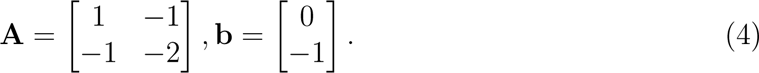

All together, the full inequality phenotype statement is,

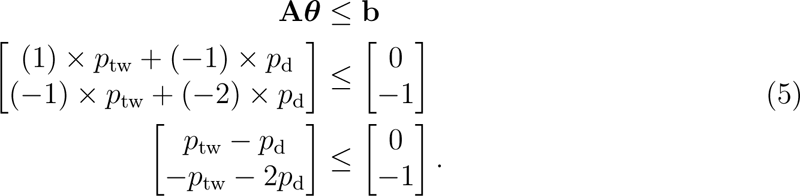

We use the same procedure to specify a number of phenotypes that together represent four complete hypothesis sets with mutually exclusive diel phenotype hypotheses: *Maximizing*, *Traditional*, *General*, and *Selection* (Figures 1, 2). The first three hypothesis sets are strictly on how much an animal uses each diel period, while the fourth (*Selection*) is based on the selection of a diel period, where the probability of use is greater than the proportional amount of time available in a diel period (Northrup et al. 2022; Gallo et al. 2022). The file Supplementary Material S2 describes and mathematically defines the set of inequalities for each phenotype.

**Figure 1:**
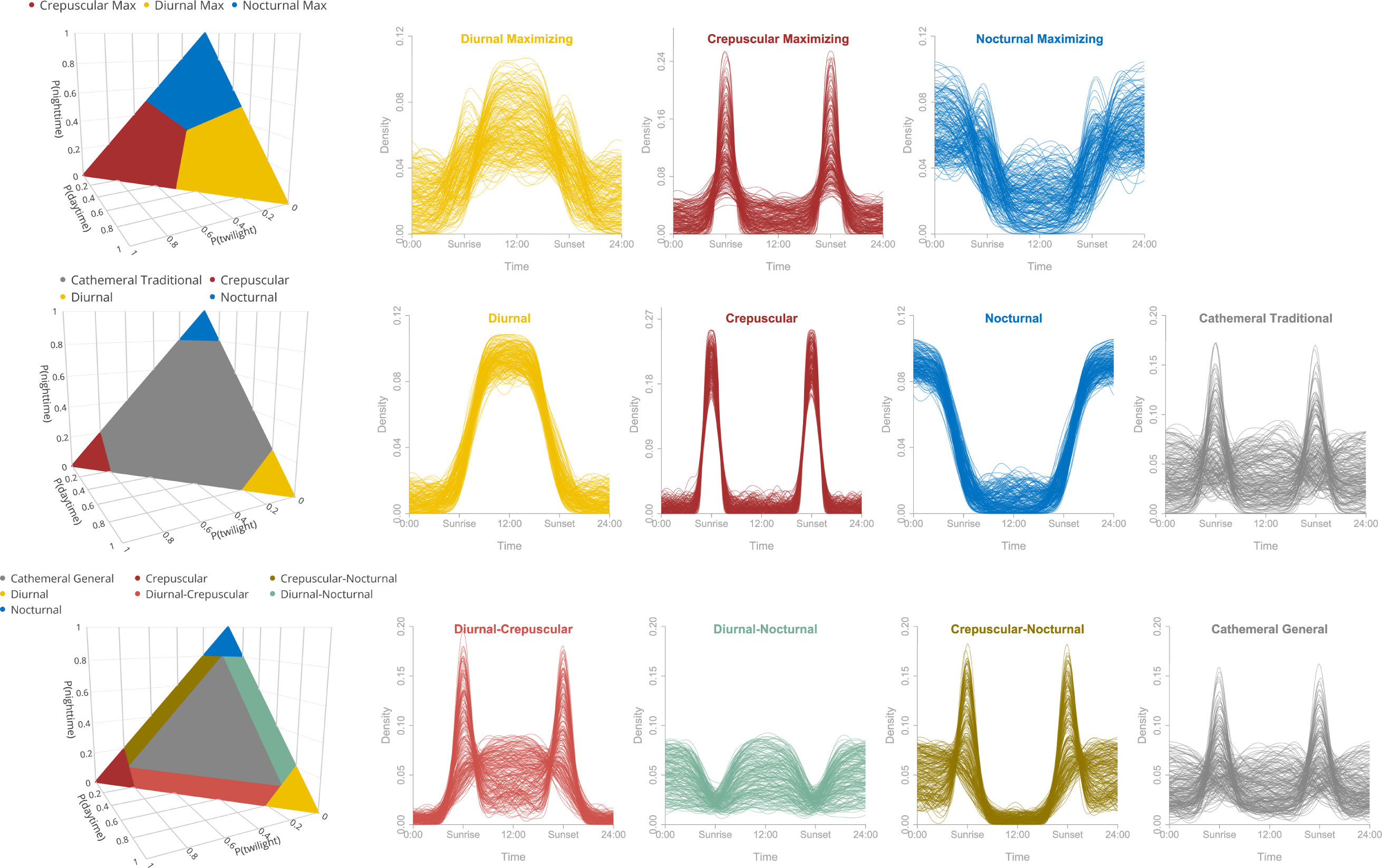
Diel niche hypothesis sets (first column, from top to bottom: Maximizing, Traditional, General) as defined by the probability of activity during daytime, nighttime, and twilight. For each hypothesis in a set, examples of circular kernel density values are plotted over the 24 hour period. Note that the General hypothesis set (3rd row) includes the diel hypotheses of diurnal, nocturnal, and crepuscular, which are plotted with the Traditional hypothesis set (second row).

### Diel Hypothesis Sets

The *Maximizing* hypothesis set includes three diel phenotype hypotheses (Figure 1; Crepuscular Max, Diurnal Max, and Nocturnal Max) with the objective of evaluating which diel period is used most. As such, there is no hypothesis about activity across multiple diel periods (i.e., cathemeral). The *Traditional* hypothesis set includes four diel phenotype hypotheses (Figure 1; Crepuscular, Diurnal, Nocturnal, Traditional Cathemeral) that aim to capture the general interpretation of these hypotheses from the literature. Crepuscular, Diurnal, and Nocturnal are defined based on having at least 0.80 probability (threshold probability, *ξ*_1_ = 0.80) in their respective diel periods. If an animal is not mostly active in one period than it is defined as Traditional Cathemeral; this occurs when either two or three diel periods are used more than 1 *− ξ*_1_. The logic behind the threshold of 0.80 is that an animal is predominately active in one diel period, but is not so strict that there can be some moderate amount of activity outside of this period. While *ξ*_1_ = 0.80 is the default value in Diel.Niche, it can easily be modified by a user when fitting these hypotheses to their data.

The *General* hypothesis set includes seven diel phenotype hypotheses (Figure 1). The Diurnal, Crepuscular, and Nocturnal hypotheses are defined the same as in *Traditional*. The main difference is the separation of the probability space of Traditional Cathemeral into four more specific hypotheses: General Cathemeral, Crepuscular-Nocturnal, Diurnal-Nocturnal, and Diurnal-Crepuscular. The General Cathemeral hypothesis—which represents a subset of the parameter space taken by the previously mentioned Traditional Cathemeral—aims to define when an animal uses all three diel periods at equal to or more than a minimum amount (i.e., *ξ*_2_ *≤ p*_tw_*, p*_d_*, p*_n_ *≤ ξ*_1_). We defined the lower threshold probability as *ξ*_2_ = 0.10, such that we consider it important to differentiate animal activity when a diel period is used at least this much (Figure 1). However, if only two diel periods are used above *ξ*_2_ then we classify this activity using one of the binomial hypotheses (Crepuscular-Nocturnal, Diurnal-Nocturnal, and Diurnal-Crepuscular). For example, suppose a species is active mainly during the day (*p*_d_ = 0.78), but is also relatively active during twilight (*p*_tw_ = 0.16), and not very active at night (*p*_n_ = 0.06). We would define this activity as Diurnal-Crepuscular because *p*_tw_, *p*_d_ *≥ ξ*_2_, while *p*_n_ *< ξ*_2_. However, if night activity was also higher than *ξ*_2_, such that *p*_d_ = 0.70 and *p*_tw_*, p*_n_ = 0.15 then we classify this activity as General Cathemeral because a moderate to large amount of activity is occurring in all three diel periods. In summary, the *Traditional* hypothesis set distinguishes between unimodal and multimodal diel activity, while the *General* set distinguishes among unimodal, bimodal, and trimodal activity.

The *Selection* hypothesis set includes seven diel phenotype hypotheses (Figure 2; Day, Day-Night, Day-Twilight, Equal, Night, Night-Twilight, and Twilight Selection). Each are defined based on an inputted (i.e., not estimated) amount of proportional time available to an animal in each diel period (**p**_avail_ = [*p*_av.tw_*, p*_av.d_]; the available time in the night period is derived as, 1 *− p*_av.tw_ *− p*_av.d_). As such, the combination of parameter values defined as a certain phenotype and the parameter space area of each phenotype will change based on **p**_avail_. The values of **p**_avail_ will depend on the day of the year and location of sampling. Each phenotype is defined based on a single diel period or multiple diel periods being used greater than available. For example, diurnal selection occurs when 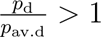 and 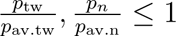. The Equal selection hypothesis is different in that it tests the equality among the probabilities and **p**_avail_.

**Figure 2:**
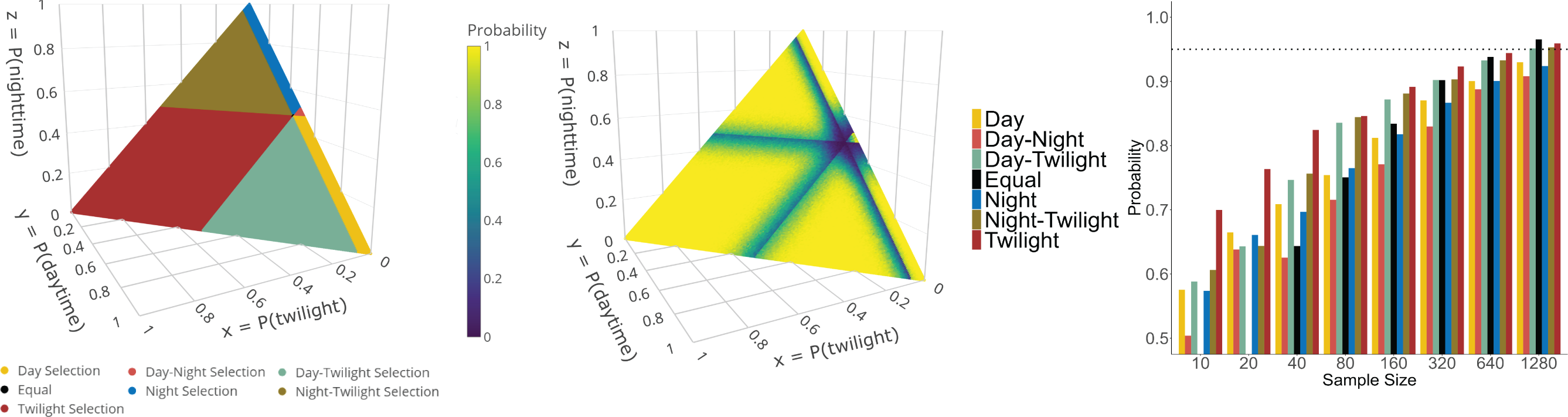
The Selection hypotheses (left) as defined by the disproportional amount of activity during daytime, nighttime, and twilight, given their respective available amount of time in each period (availability defined as 0.04, 0.48, and 0.48 for the respective diel periods). The middle plot depicts the probability that the generating model is most supported by parameter combinations when the sample size is 100 total observations. The right plot comprises results of the probability the generating model was most supported across varying sample sizes. Complementary simulation results are found in Figure S4.

We offer these definitions as a means to standardize the diel activity hypothesis language. Each hypothesis set is preset and available for use in Diel.Niche. The function ‘hyp.sets’ can be used to list the code names for each hypothesis and can be used to pass to the ‘triplot’ function to provide a 3D plot of the parameter space (Figure 1; first column) as,

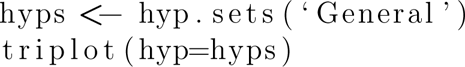

However, the package also allows the specification of new hypotheses (see ‘Additional Resources’). Researchers interested in changing the probability threshold values (*ξ*_1_ and *ξ*_2_) or defining novel hypotheses should report these in their research.

### Modeling and Estimation

Applying inequality constraints on the parameters of multinomial models is a well known modeling and estimation issue in statistics (Silvapulle & Sen, 2011). Implementing these models is often challenging in general-purpose software. We used a Bayesian approach and take advantage of a general Gibbs Markov-chain Monte Carlo (MCMC) sampler developed and implemented in the R package multinomineq (Heck et al., 2019). The sampler is highly efficient and implemented in C++, making the MCMC sampler relatively fast compared to general-purpose software implementations. Generalized model specification and the likelihood function are fully detailed in Heck et al. 2019. The core modeling and estimation in the Diel.Niche package are done using wrapper functions that implement more general functions of the multinomineq package. In terms of model parameter (***θ***) prior specification, probabilities are given a Dirichlet prior distribution with a truncated support that is defined by the constrained parameter space (Heck et al., 2019). The default hyperparameters used in Diel.Niche for the prior are shape parameters of 1 for each free model parameter. This is equivalent to a multi-parameter uniform prior for each parameter, which is thus a very uninformative prior.

Model comparison is done via Bayes factors (Berger, 2013) for models with inequality constraints (Klugkist & Hoijtink, 2007), which implicitly provides a trade-off between model fit and complexity. Each model within a hypothesis set requires a prior to compute posterior model probabilities from the Bayes factors. Diel.Niche uses a default of equal weight among the set, but this can be changed. For example, if the literature attributes a species as nocturnal, a researcher may want to increase the prior of the nocturnal hypothesis. The prior will therefore depend on the goal of the study.

## Things to consider before using this method

### The Data

The study of diel activity describes when behavioral actions occur within the 24-hour light-dark cycle. However, capturing all animal behaviors is generally impossible without direct and continuous behavioral sampling. This is often not possible for many species that cannot be followed to record behavioral observations without changes in animal behavior occurring due to human presence. Even when individual animals can be followed, tracking continuously over many 24-hour periods is simply logistically infeasible. Here, we focus on the traditional definition of diel activity in terms of animal movement (Hut et al., 2012) as it is a fundamental behavior to ecosystem functioning and species survival (Tucker et al., 2018). Increasingly, high frequency GPS telemetry is used to sample the location and movement of many species. These datasets could be very useful to estimate diel activity, especially as behavioral state modeling may help separate when and where specific behaviors are occurring (Michelot and Blackwell, 2019). Limitations of these data include the 1) expense in tracking many individuals in a population to gain a population-level inference, 2) difficulty in trapping many rare or small species, which are not readily or legally able to be trapped, and 3) morphology of some species which poses challenges for external attachment (e.g., neck collars) thereby necessitating possibly risky surgery for a transmitter to be implanted.

The type of data that primarily motivated this work was species detections from trail cameras. However, any spatially-fixed technology that samples species activity throughout the 24-hour period continuously is applicable. This could include autonomous audio recording units, where species can be identified and calling conveys an important activity. The fundamental data unit we are considering is a set of detection frequencies (**y**). Considering common trail camera sampling designs, where a set of *n* cameras are deployed at unique spatial locations over a period of interest (e.g., a climatic season), **y** may be an aggregation of detections across all cameras for this specified period of time. Thus, inference is at the study area and sampling period level. However, if enough data are available inference could be made either at the camera trap level or among a subset of cameras within a study area (e.g., cameras placed in different habitat types).

We assume that dependencies in the data have been removed, such as temporal autocorrelation among a species’ detections at a sampling location. Thus, frequencies of detections during twilight, daytime, and nighttime are independent. For camera trap sampling, it is common to have several-to-many sequential photographs of the same individual at the same location. Commonly, correlation is removed by subsetting data to exclude detections within a specified period of time (e.g., 10, 15, 30 minutes; Farris et al. 2015). A more quantitative approach would be to test this correlation using lorelograms (Iannarilli et al., 2019).

Of fundamental importance is the classification of each independent detection into the appropriate diel period. Fortunately, this can be be determined if you know the date, time, and spatial location of a given sample. This information can be used to calculate the start and end of various sunlight phases (e.g., sunrise, sunset, dusk) and are implemented in R with the suncalc package (Thieurmel and Elmarhraoui, 2022). We suggest samples be assigned to the three diel periods as shown below.

- **Twilight:** The sample occurred during either dawn or dusk. Dawn is the time between the start of morning astronomical twilight and the end of morning nautical twilight. Dusk is the time between the start of evening nautical twilight and the end of evening astronomical twilight.
- **Daytime:** The sample occurred during the day, which is the time between the end of morning nautical twilight and the start of evening nautical twilight. Note that this time period also includes morning and evening civil twilight, which are time periods where light is available while the sun is not visible.
- **Nighttime:** The sample occurred during the night, which is either between midnight of that day and the start of morning astronomical twilight or between the end of evening astronomical twilight and midnight of the following day.

Using this classification scheme makes it possible to generate a frequency table of detections that can then be used within the Diel.Niche package.

### Hypothesis Discrimination

Prior to implementing this framework, it is important to understand how sample size affects the ability to discriminate among hypotheses within a set. For each of our four hypothesis sets, we simulated 4000 datasets under each hypothesis within a set at a wide range of total samples sizes (10, 20, 40, 80, 160, 320, 640, 1280; 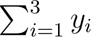). We fitted each dataset and estimated the posterior model probability of the hypothesis generating the data; each model was given equal prior weight. Since each hypothesis is mutually exclusive (i.e., non-overlapping defined parameter space), model uncertainty will be highest when the true probability parameters are near the boundary of a hypotheses parameter space and thus adjacent to another hypotheses parameter space. This will occur more often with hypotheses with small parameter spaces (e.g., *General* hypotheses). We considered how this would alter model selection and certainty by only simulating data such that hypotheses parameter spaces are separated by at least 0.1 and 0.2 (Figure S2). Lastly, we explicitly demonstrate how the arrangement of hypotheses’ parameter space affects the ability to identify the hypothesis generating the data. To do so, we use the *Selection* hypothesis set to simulate 200 datasets with a sample size of 100 for each combination of parameter values (***θ***) at increments of 0.05 and estimate the probability the generating hypothesis is most supported within the set. We set *p*_avail_ = [0.04 0.48].

We found the ability to identify the generating model with the most support (i.e., highest posterior model probability) to vary by hypothesis set (Figures 2, 3, S3, and S4). Considering the *Maximizing* hypothesis set, 320 independent samples were required to obtain a 95% probability that the most supported model in the set was the generating model. There was little variation by hypothesis because all hypotheses had the same area of parameter space and length of parameter space adjacent to another hypothesis. We also found that when separating hypotheses by a probability of 0.1 and 0.2, only 40 and 20 samples, respectively, were needed to obtain a 95% probability. Thus, as probabilities are further from equal (*p*_tw_ = *p*_d_ = *p*_n_ = 0.333), the easier it is to discriminate among the hypotheses in the *Maximizing* set with high probability.

Considering the *Traditional* hypothesis set, 10 independent samples were needed to obtain a 95% probability that the most supported model in the set was the generating model for the Crepuscular, Diurnal, and Nocturnal hypotheses (Figures 3 and Figures S3). There was considerably more model uncertainty for the Traditional Cathemeral hypothesis. This hypothesis required 640 samples to achieve the same probability. However, when separating hypotheses by a probability of 0.1 and 0.2, only 80 and 40 samples were required, respectively. Thus, in cases where Traditional Cathemeral hypothesis probabilities are more equal, less data is needed to identify the data generating hypothesis with high probability.

**Figure 3:**
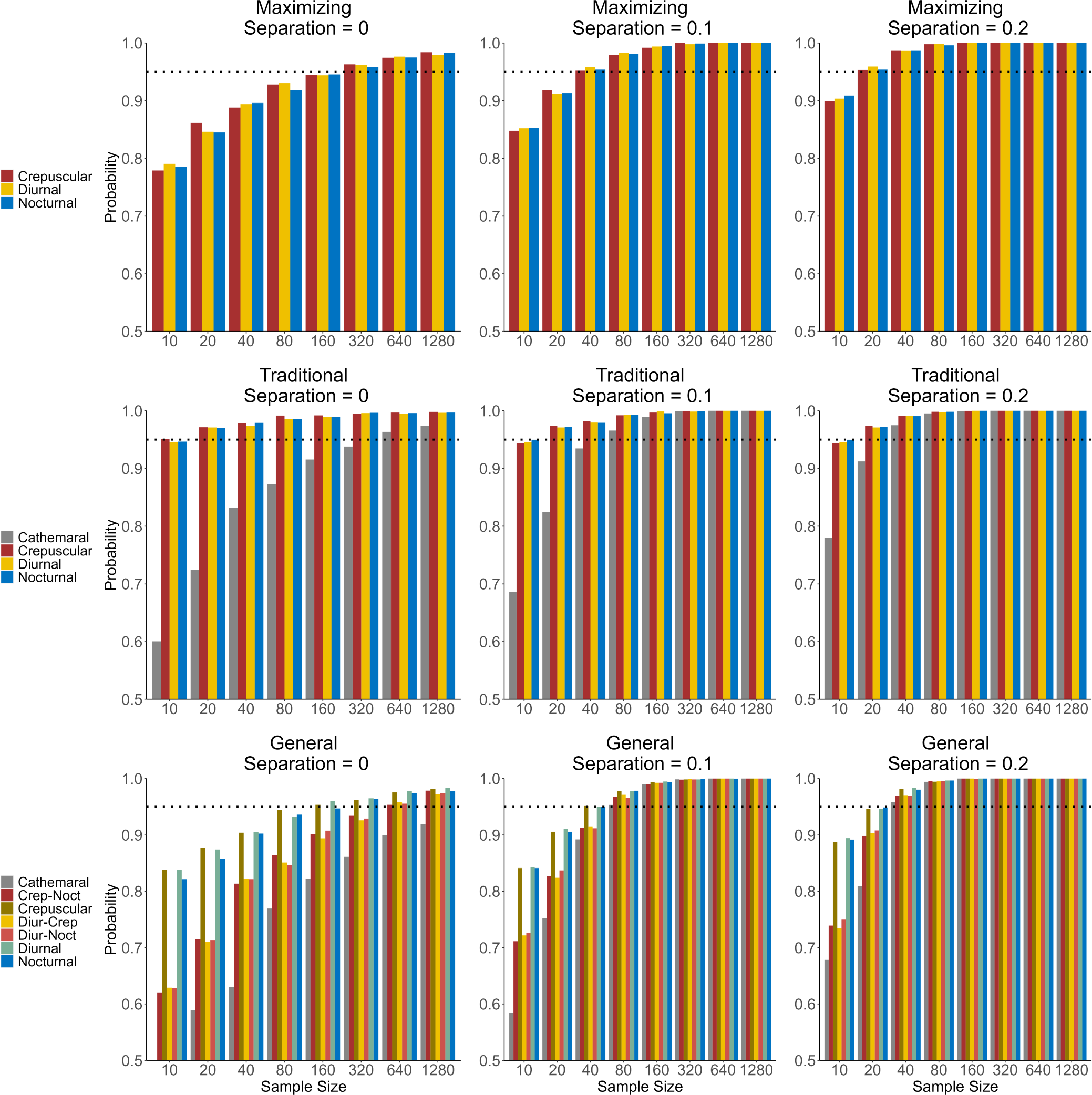
Simulation results of the probability the generating model was most supported across varying sample sizes and the extent of separation in probability space between hypotheses within a hypothesis set. Complementary simulation results are found in Figure S3.

Considering the *General* hypothesis set, 160 independent samples were required to obtain a 95% probability that the most supported model in the set was the generating model for the Crepuscular, Diurnal, and Nocturnal hypotheses (Figures 3 and Figures S3). The bimodal hypotheses required 640 samples, while 1280 samples were still not enough for the Cathemeral General hypothesis to obtain a 95% probability of support. Again, separating the hypotheses increases the ability to discriminate among them. Since these hypotheses have less parameter space and more adjacency to other hypotheses parameter space, discriminating among them will require larger sample sizes compared to hypotheses of the other sets. For the Cathemeral General hypothesis, there is also higher implicit variance in the probabilities as they get closer to 0.5, which also makes it harder to discriminate against the other hypotheses.

Considering the *Selection* hypothesis set, 1280 independent samples or more were required to obtain a 95% probability that the most supported model in the set was the generating model (Figures 2 and Figures S4). The twilight selection hypothesis was more easily identified as the generating model for smaller sample sizes because it has the largest parameter space area. These results would suggest this hypothesis set to be only moderately useful to many realistic sample sizes. However, looking at how the probability of the generating model being most supported varies throughout the parameter space, we see a large set of parameter values where it is easy to identify the generating model (probability near 1) at a sample size of 100 (Figure 2). The ability to identify the generating model in this hypothesis set and all others depends strongly on how close the true probabilities are to one or more other hypotheses, which depends on the area of parameter space for the hypothesis (Figure S5).

### The Unconstrained Model

What is the benefit of constraining model parameters? In other words, why not fit an unconstrained multinomial model, which can more easily be specified hierarchically to link probabilities of activity to spatial/temporal covariates (see Gallo et al. 2022)? The main reason is that our aim is to focus on *a priori* definitions of diel phenotypes, so that we can offer them as hypotheses to be evaluated using empirical data. By constraining parameters, we can focus on the hypothesis and not simply a change in probabilities. Considering inference across data analysis units (multiple **y** values pertaining to different study areas or sampling periods) or studies allows an accurate interpretation of diel niche switching. Second, for a given hypothesis, model parameters will be more precise because of the constraints. Thus, even with small sample sizes parameter estimates will be more precise compared to a model with unconstrained parameters. Lastly, we do not need to ignore the unconstrained model and can actually incorporate it into our hypotheses sets in the Diel.Niche package (see Additional Resources).

For researchers who take an alternative procedure to model diel activity, we encourage them to consider post hoc classification of their inference using the diel phenotypes defined here. If results can be summarized into the three probabilities (*p*_tw_, *p*_d_, *p*_n_) then this framework could be of use (see Worked Examples). Specifically, it may help differentiate between changes in activity within a diel phenotype and the switching to a new diel phenotype (e.g., nocturnal → diurnal).

## Worked Examples

The fundamental data analysis unit (**y**) should be considered as a set of observations of a species of interest that was continuously sampled throughout the 24-hr period in a place and time of interest. In the following, we consider four worked examples with varying goals that demonstrate the use of the Diel.Niche package. The first three examples are based on camera trap data collected at 131 spatial locations deployed through urban greenspace in Chicago (Illinois, USA) to better understand the spatio-temporal dynamics of urban mammals (Magle et al., 2019).

### Single data unit analysis

Here, we are interested in classifying the traditional diel niche of the Virginia opossum (*Didelphis virginiana*) in an urban environment during the winter of 2018-2019. Specifically, sampling occurred for a total of 27 days from December 29*^th^* to January 24*^th^*. These data are available in the package using the ‘diel.data’ object. After removing sequential photos at the same site within a 15 minute period, we aggregated these observations across all camera locations, such that **y**= [5 20 30]. A vignette describing the full set of code for this analysis can be found on GitHub and as part of the package.

Using the ‘diel.fit’ function to estimate model probabilities assuming equal weights on each model,

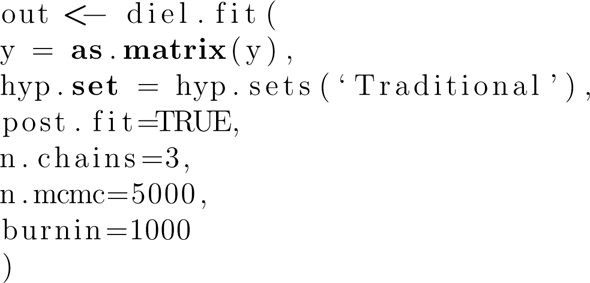

we find that the cathemeral hypothesis is supported with a posterior model probability of 1.0. Using base R plotting and the ‘triplot’ function as triplot(out) we can visualize the posterior distributions of the activity probabilities (Figure 4).

**Figure 4:**
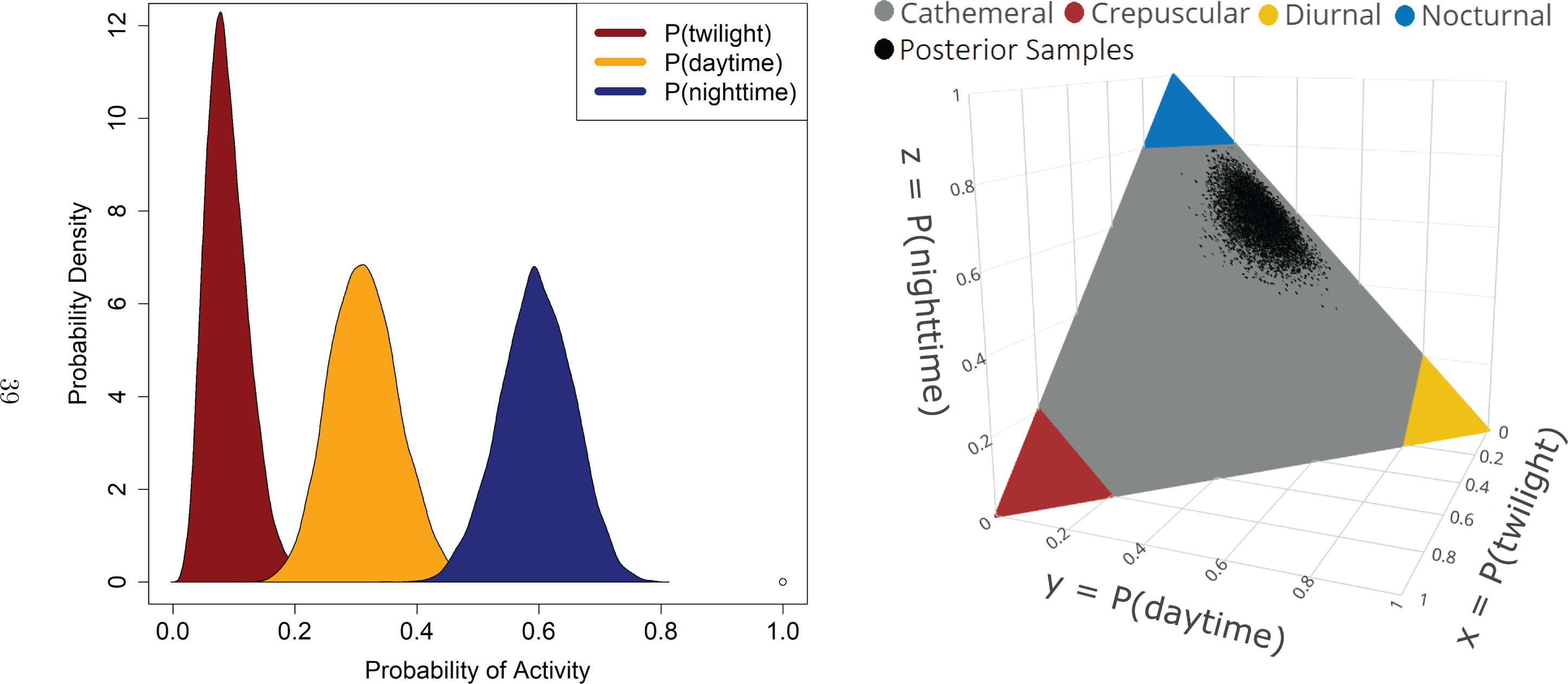
Posterior distributions of the probability of activity (left) and the same in three-dimensional form (right) overlaying the Traditional hypothesis set for the Virginia opossum sampled in Chicago, IL USA in the winter of 2018.

We see the opossum is most active at night in the winter, but also has a relatively high amount of activity during the daytime. The 3D figure allows us to consider where in the *Traditional* hypothesis set these probabilities fall. Given the low amount of activity at twilight, we find using the *General* hypothesis set that the Diurnal-Nocturnal hypothesis would be a more accurate description of the opossum’s winter diel niche.

### Multiple species with one data analysis unit

Commonly with passive sampling techniques, such as trail cameras, the focus of a study is on a community rather than a single species. Here, our objective is to evaluate the diel niche of a community of urban mammals in the winter by considering support for hypotheses from the *General* hypothesis set. Again, we use the ‘diel.data’ object to obtain the aggregated data from trail cameras deployed in Chicago (Illinois, USA) for 25 days between January 25*^th^*, 2018 and February 20*^th^*, 2019. We estimated and compared the diel niche of each observed species following the same procedure as in the first example. A vignette describing the full set of code for this analysis can be found on GitHub and as part of the package.

A total of seven native terrestrial mammal species were observed with total species detections ranging from 25-284 (Table 1). We found a high-level of evidence (model probability *>* 0.9) that white-tailed deer (*Odocoileus virginianus*) are Diurnal-Nocturnal, northern raccoon (*Procyon lotor*) and eastern cottontail (*Sylvilagus floridanus*) are Nocturnal, and the gray squirrel (*Sciurus carolinensis*) and fox squirrel (*Sciurus niger*) are Diurnal. We are moderately confident (model probability near 0.75) that coyote (*Canis latrans*) and Virginia opossum (*Didelphis virginiana*) are Diurnal-Nocturnal. Comparing these results to the literature (Cox et al. 2021, Supplementary Data 2), we see agreement with only the northern raccoon and both squirrel species. There is disagreement with all other species, where the literature considered the diel niche of the Virginia opossum to be nocturnal, and the white-tailed deer and the eastern cottontail to be crepuscular. The coyote diel niche difference is more subtle, where they are considered as cathemeral in the literature, we estimated their activity to be a specific form of cathemerality (i.e., Diurnal-Nocturnal).

**Table 1:** Detection frequencies of urban mammals in Chicago, Illinois USA during the winter of 2019 and posterior model probabilities of support from the General hypothesis set. Diur = Diurnal, Noct = Nocturnal, and Crep = Crepuscular

| Species | Twilight | Day | Night | Diurnal | Nocturnal | Crepuscular | Cathemeral | Diur-Crep | Diur-Noct | Crep-Noct |
| --- | --- | --- | --- | --- | --- | --- | --- | --- | --- | --- |
| Coyote | 3 | 8 | 23 | 0.00 | 0.07 | 0.00 | 0.17 | 0.00 | 0.75 | 0.00 |
| Virginia Opossum | 0 | 7 | 18 | 0.00 | 0.21 | 0.00 | 0.02 | 0.00 | 0.77 | 0.00 |
| White-tailed Deer | 4 | 39 | 20 | 0.00 | 0.00 | 0.00 | 0.07 | 0.00 | 0.93 | 0.00 |
| Northern Raccoon | 6 | 8 | 133 | 0.00 | 1.00 | 0.00 | 0.00 | 0.00 | 0.00 | 0.00 |
| Eastern Gray Squirrel | 0 | 284 | 0 | 1.00 | 0.00 | 0.00 | 0.00 | 0.00 | 0.00 | 0.00 |
| Eastern Fox Squirrel | 0 | 49 | 0 | 1.00 | 0.00 | 0.00 | 0.00 | 0.00 | 0.00 | 0.00 |
| Eastern Cottontail | 13 | 5 | 123 | 0.00 | 0.99 | 0.00 | 0.00 | 0.00 | 0.00 | 0.01 |

Using the most supported model by species, we plotted the estimated probability of diel activity for the three diel periods (Figure 5). Interestingly, the coyote is active in all three diel periods, but is most active during the night, which is also when their prey species, the eastern cottontail, is active. White-tailed deer were found to be most active during the daytime, which is surprising given that they are not avoiding periods of high human activity in an urban environment. We also found that the eastern cottontail has the most amount of activity during the twilight from this mammal community.

**Figure 5:**
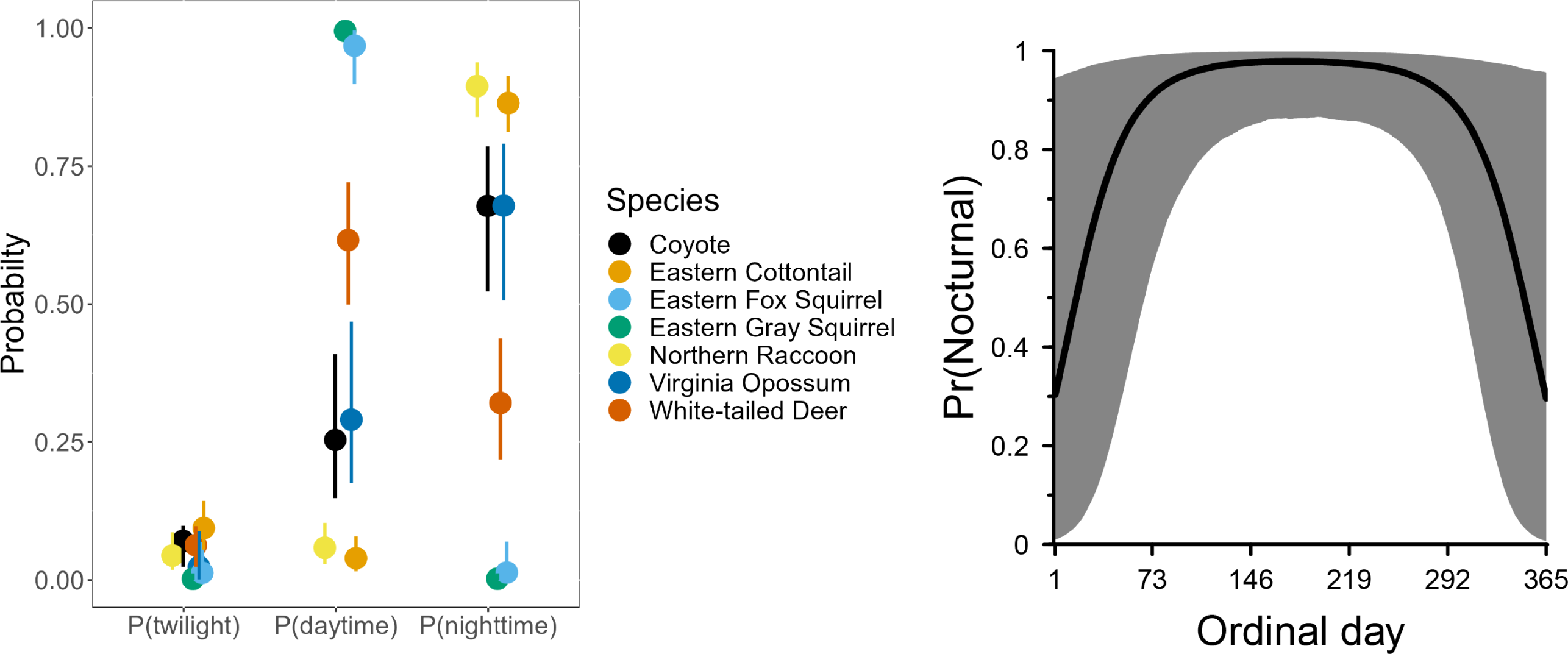
The left subplot comprises results from worked example 2: the posterior median (circles) and 95% credible intervals of the probability of activity for the mammal community sampled in Chicago, IL USA in the winter of 2019; parameters are from the most supported model for each species using the General hypothesis set. The right subplot is a result from worked example 3: the posterior predicted probability of the nocturnal hypothesis for the Virginia opossum in Chicago, IL USA.

### One or more species with multiple data units

Our framework has outlined a process for making inference on the diel niche of species for a given spatial area and sampling period. A natural question is how to link hypotheses about how the diel niche of a species may change over time or over space. Simply, how do we link spatial or temporal covariates to findings of diel niche classification? Ideally, a fully hierarchical model that synthetically connects the diel niche classification to covariates would be specified, such that parameter estimation is done jointly and all parametric uncertainties are fully recognized. However, the constrained optimization of the multinomial model makes this challenging at this time.

We offer a two-staged modeling process that accomplishes the goal. First, estimate the diel niche for a species at each spatial or temporal period of interest to identify the most supported hypothesis. Second, use the hypotheses as data in a subsequent categorical regression model that links the hypotheses to covariates of interest. It should be understood that if there is uncertainty in the most supported model then this does not get acknowledged in the regression model. To remediate this concern, researchers may want to only consider using hypotheses with a given high level of probability of support (e.g., model probability *≥* 0.80) or propagate this uncertainty into the subsequent analysis.

As one example, Virginia opossum are a predominately nocturnal species that become more active during the day in the winter in temperate environments (Gallo et al. 2022). As a result, we might expect this species to switch between diel phenotypes over the course of a year. As the previous Chicago data is from a long-term biodiversity monitoring survey, we compiled 27 different sampling units, each of which represented roughly 28 days of sampling across 131 sites in Chicago, Illinois. To capture seasonal variation, cameras were deployed in January, April, July, and October, and data comes from between 2013 and 2019 (all of which is available in the ‘diel.data’ object).

We fitted a *Traditional* hypothesis set to each analysis unit with equal prior weights on each model, and retained each analysis unit that had a probability of model support *≥* 0.80. This resulted in a total of 23 analysis units. We then fitted a categorical regression model to these data. More specifically, we let *m_i_* be a numeric representation of the diel phenotype for the i*^th^* analysis unit, such that 1 = Cathemeral, 2 = Diurnal, and 3 = Nocturnal. While there is the possibility of there being 4 diel categories, in this specific case opossum were either classified as nocturnal or cathemeral. As a result, we dropped out the crepuscular category, but retained the diurnal category to demonstrate the use of this model across more than two diel phenotypes. Further, let ***p*** be a vector of probabilities that sum to 1 and represent the probability a given diel phenotype is used. The most basic model without covariates is then

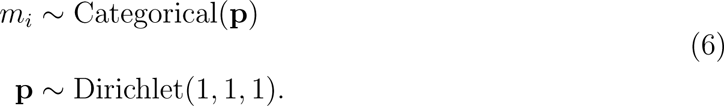

To expand the model, we connect **p** with a vector of covariates that varies across sampling units by using the softmax function, treating cathemeral as the reference category. For this analysis, we consider whether the time of year affects the opossums diel niche. Specifically, we used the average ordinal date of an analysis unit as well as the square of that ordinal date (to allow for a non-linear relationship) as covariates, such that

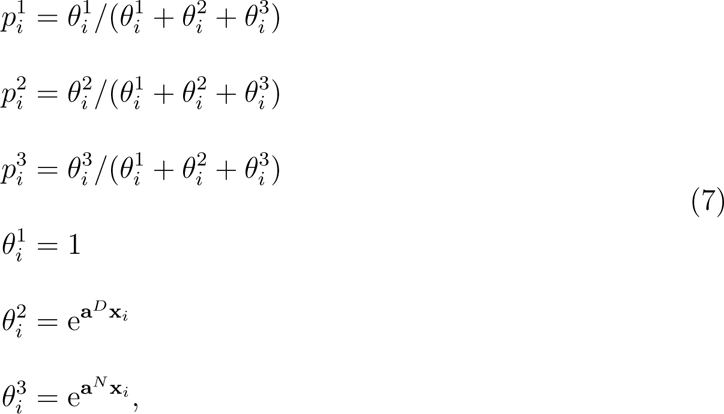

where **a***^D^* and **a***^N^* are a vector of conformable regression coefficients for diurnal and nocturnal categories while **x***_i_* is a vector of covariates with the first element being a 1 for the intercept. Weakly informative priors were used for all regression coefficients as Normal(0, 2). A vignette describing the full set of code for this analysis can be found on GitHub and as part of the package.

We fitted this model in v0.12.2 of Nimble (de Valpine et al., 2017) and used slice samplers for all parameters. Following a 10,000 step burnin we sampled parameter posterior distributions for a total of 80,0000 iterations across 4 chains. Overall, we found evidence that the probability Virginia opossum were nocturnal greatly decreased at the start and end of the year, when it is coldest in Chicago (Figure 5). This pattern was largely determined by the squared ordinal day term, which was strongly negative (*a^N^* = −1.36, 95% CI = −2.61, −0.27). This switch in diel phenotypes was quite dramatic, with the probability of nocturnality ranging from a high of 0.98 (95% CI = 0.86, 1.00) on June 6th to a low of 0.30 (95% CI = 0.01, 0.96) on December 31st.

### Diel classification with a kernel density analysis

A common statistical approach used in the diel ecology literature is that of the non-parametric circular kernel density estimators (Ridout and Linkie 2009; Figure 6). Researchers can complement this analysis type by using the Diel.Niche package to define their findings in terms of diel phenotypes. After using the overlap R package’s ‘densityPlot’ function to fit the kernel density estimator, the outputted object (kernel.out) can be passed to the Diel.Niche function ‘prob.overlap’ to integrate under the curve at the intervals along the x-axis that correspond to the diel periods (twilight, daytime, nighttime) to estimate these probabilities as,

**Figure 6:**
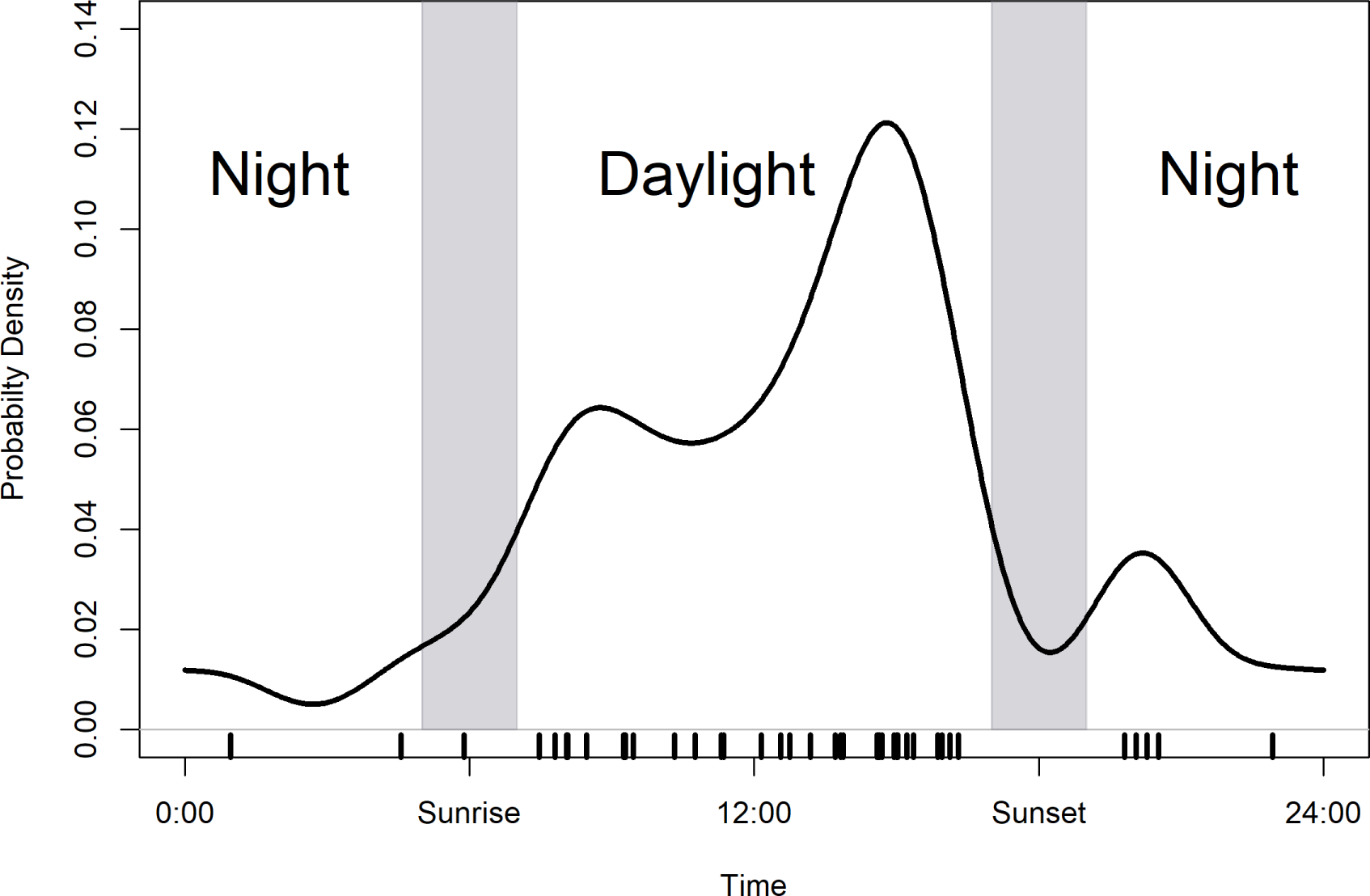
Circular kernel density estimate of an example dataset. The shaded gray area indicates periods of twilight and the tick marks along the x-axis indicate the time of each observation. Using the hypothesis sets of Maximizing, Traditional, and General lead to classifying this activity pattern as the hypotheses Diurnal Max, Crepuscular Traditional, and Diurnal-Nocturnal, respectively.

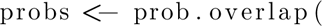

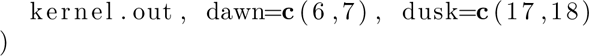

The other input required are the numeric values defining the beginning and end periods of dawn and dusk. Using example data from Figure 6, the probabilities of twilight, daytime, and nighttime were estimated as 0.055, 0.748, and 0.194, respectively. From here, the probabilities can be passed to the function ‘posthoc.niche’ that will match the diel phenotype within a hypothesis set that includes these probabilities as,

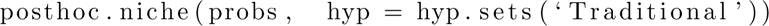

Using the *Traditional* hypothesis set, the results depicted in Figure 6 would be defined as the Cathemeral Traditional hypothesis. This may be unintuitive simply by looking at the plot. While there is a large amount of activity during the daytime, the probability of nighttime use is also quite high at a probability of 0.194, thus supporting the Cathemeral designation. If we use the *Maximizing* hypothesis set, we find that the Diurnal Max hypothesis is most supported, while using the *General* hypothesis set, the most supported hypothesis is more specifically the Diurnal-Nocturnal hypothesis. A vignette describing the full set of code for this analysis can be found on GitHub and as part of the package.

By defining the diel hypotheses *a priori* and explicitly, we can more accurately make inference based on a given objective. Visually, the kernel density plot shows this animal is most active during the daytime, which is confirmed with the Diurnal Max hypothesis being most supported in the *Maximizing* hypothesis set. However, considering more traditional definitions of diel activity, we should define this animal as cathemeral, given its support under the Cathemeral *Traditional* hypothesis. More so, if we want to make a clear delineation between activity at two diel periods versus three diel periods, the *General* hypothesis set makes it evident the species is mostly active during only two diel periods, as supported by the Diurnal-Nocturnal hypothesis.

## Caveats

Our goal was to focus on defining, estimating, and modeling hypotheses of diel phenotypes motivated by the animal ecology and evolution literature. It is important to recognize the probability of activity in the three diel periods is a simplification of a complex process (e.g., see kernel density plots in Figure 1; Gaston 2019; Gilbert et al. 2022). More detailed inference, how animals use time from minute to minute or hour to hour are warranted and needed. However, we hope that researchers will consider how they translate their findings into standardized diel language and hope that the Diel.Niche package will be of use in doing so.

For three of the hypotheses sets (*Maximizing*, *Traditional*, *General*) the focus is on characterizing an animal’s diel phenotype based on activity use in a diel period. It is important to understand the equivalence of how we defined diurnal, crepuscular, and nocturnal. Meaning that each is defined with the same constraints of how much activity is required to occur in their respective diel periods, regardless of the amount of time available to the animal in each period. We think this equivalence is important and appropriate for these diel phenotypes to have the same meaning in characterizing when the animal is most active. However, this has especially important implications for identifying an animal as crepuscular in these hypotheses sets.

Given our definition for the twilight period there will only ever be a small amount of time available for the animal (e.g., *<* 0.10), such that it may be unusual to find an animal using the twilight with a probability of 0.80 or greater (the lower constraint for defining crepuscular in the *Traditional* and *General* hypothesis sets). One option would be to modify our definitions of twilight and increase the amount of time defined as twilight when classifying observations. Rather than strictly defining twilight around sunlight times, which leaves only small portion of the day available for a species to be crepuscular, there may be biological reasons to expand twilight to encompass some part of the end of night and the beginning of day. However defined this should be documented when reporting results. It is also important to consider when using data from the northern and southern latitudes during the summer and winter months there are extremes in the amount of time available in each diel period. However, if the inference being sought is on how animals use each diel period, given the amount of time available to them, this requires focusing on selection of a given diel period. As such, the *Selection* hypothesis set can be used to complement inference from the other hypotheses sets.

## Additional Resources

The main vignette for the package Diel.Niche is provided on the GitHub repository demonstrating key functions. This includes simulating data under a hypothesis, model fitting and estimation, plotting, and example data. This vignette can also be accessed in the package (assuming vignettes have been built during package installation) as vignette(‘Diel-Niche-Main’). A secondary vignette (‘Diel-Niche-Additional’) is also available that describes additional hypotheses (defined and described in Supplementary Material S2), as well as how a user may change default values (e.g., *ξ*_1_ and *ξ*_2_) and even specify novel hypotheses. Further, each worked example is provided as a vignette in the package or on Github, as well as a fifth worked example demonstrating how to make inference from the *Selection* hypothesis set.

Users who prefer a graphical interface can use Diel.Niche through the R Shiny web application (https://shiny.celsrs.uri.edu/bgerber/DielNiche/). The current implementation allows users to specify a single data analysis unit **y** and to choose among the four hypotheses sets. The application will automatically generate a table of model probabilities and posterior quantities of model parameters. Both a 2D and 3D plot of model parameters are generated and can be downloaded.

## Conclusion

Studies of animal diel ecology and evolution provide inference to how animal activity is shaped within the 24-hour light-dark cycle. This leads to a more complete understanding of an animal’s niche (Cox et al., 2021), as well as having implications for how to conserve wild animal populations (Rivera et al., 2022; Cox et al., 2023b). As non-invasive sampling and animal-borne sensors continue to evolve and make it easier to continuously collect data on wild animals throughout the diel cycle, we expect increasing studies focused on how animals use diel time.

We offer the Diel.Niche R package as a framework to quantitatively define and thus standardize diel ecology language regarding diel phenotypes. We hope this will lead to more accurate comparisons across studies and reduce potential confounding from qualitative and visual interpretations of diel phenotypes. We offer several complete diel phenotype hypotheses sets that aim to accomplish different study objectives. The Diel.Niche package can be used to compare models, estimate parameters, and visualize results, or be used to complement other analyses by defining their inference into standardized terminology. We suggest researchers report their results following similar text as – “Using the *<*insert name*>* hypothesis set, we found that *<*species name*>* were *<*diel phenotype*>* (p = *<*insert model probability*>*), spending *x* (95% CI = low, hi) amount of time active during the night, *y* (95% CI = low, hi) amount during the day, and *z* (95% CI = low, hi) amount during twilight.” Ultimately, we hope our framework and R package will assist resource managers and researchers to more clearly and effectively define and differentiate how animals use diel time to improve conservation approaches.

## Author Contributions

Journal Submission and Manuscript Type: Journal Animal Ecology, Research Methods Guides

*These are instructional papers that aim to serve as a practical guide for animal ecologists in using a specific experimental or theoretical model or system, or software package. Research Methods Guides tend to be no longer than 7000 words (including title page, abstract, and references list)*.

## Acknowledgments

We are grateful to the 200+ data contributors to the Global Animal Diel Activity Project (https://diel-project.github.io/) that motivated this manuscript and the Diel.Niche R package. We are also grateful to the Urban Wildlife Information Network, coordinated by Lincoln Park Zoo’s Urban Wildlife Institute, for the use of data within the package and as part of the worked examples. This work was supported by the USDA National Institute of Food and Agriculture, Hatch Formula Project 1017848.

## Data Availability Statement

All data are available in the R package on GitHub at (https://github.com/diel-project/Diel-Niche-Modeling).

## References

Anderson, S. R., & Wiens, J. J. (2017). Out of the dark: 350 million year‘s of conservatism and evolution in diel activity patterns in vertebrates. Evolution, 71, 1944–1959. f

Berger, J. O. (2013). Statistical decision theory and Bayesian analysis. Springer Science & Business Media.

Cox, D. T. C., Gardner, A. S., & Gaston, K. J. (2021). Diel niche variation in mammals associated with expanded trait space. Nature communications, 12, 1753.

Cox, D. T., Baker, D. J., Gardner, A. S., & Gaston, K. J. (2023). Global variation in unique and redundant mammal functional diversity across the daily cycle. Journal of Biogeography, 50, 629–640.

Cox, D. T., Gardner, A. S., & Gaston, K. J. (2023). Diel niche variation in mammalian declines in the Anthropocene. Scientific Reports, 13, 1031.

Cunningham, C. X., Scoleri, V., Johnson, C. N., Barmuta, L. A., & Jones, M. E. (2019). Temporal partitioning of activity: Rising and falling top-predator abundance triggers community-wide shifts in diel activity. Ecography, 42, 2157–2168.

de Valpine, P., Turek, D., Paciorek, C. J., Anderson-Bergman, C., Lang, D. T., & Bodik, R. (2017). Programming with models: writing statistical algorithms for general model structures with NIMBLE. Journal of Computational and Graphical Statistics, 26, 403–413.

Farris, Z. J., Gerber, B. D., Karpanty, S., Murphy, A., Andrianjakarivelo, V., Ratelolahy, F., & Kelly, M. J. (2015). When carnivores roam: temporal patterns and overlap among M adagascar’s native and exotic carnivores. Journal of Zoology, 296, 45–57.

Frey, S., Fisher, J. T., Burton, A. C., & Volpe, J. P. (2017). Investigating animal activity patterns and temporal niche partitioning using camera-trap data: Challenges and opportunities. Remote Sensing in Ecology and Conservation, 3, 123–132.

Gaston, K. J. (2019). Nighttime ecology: the “nocturnal problem” revisited. The American Naturalist, 193, 481–502.

Gaynor, K. M., Hojnowski, C. E., Carter, N. H., and Brashares, J. S. (2018). The influence of human disturbance on wildlife nocturnality. Science, 360, 1232–1235.

Gallo, T., Fidino, M., Gerber, B., Ahlers, A. A., Angstmann, J. L., Amaya, M., et al. (2022). Mammals adjust diel activity across gradients of urbanization. Elife, 11, e74756.

Gilbert, N. A., McGinn, K. A., Nunes, L. A., Shipley, A. A., Bernath-Plaisted, J., Clare, J. D., … & Zuckerberg, B. (2022). Daily activity timing in the Anthropocene. Trends in Ecology & Evolution, 38, 324–336.

Hall, M. I., Kamilar, J. M., & Kirk, E. C. (2012). Eye shape and the nocturnal bottleneck of mammals. Proceedings of the Royal Society B: Biological Sciences, 279, 4962–4968.

Heck, D. W., & Davis-Stober, C. P. (2019). Multinomial models with linear inequality constraints: Overview and improvements of computational methods for Bayesian inference. Journal of mathematical psychology, 91, 70–87.

Hut, R. A., Kronfeld-Schor, N., van der Vinne, V., & De la Iglesia, H. (2012). In search of a temporal niche: environmental factors. Progress in Brain Research, 199, 281–304.

Hutchinson, G. E. (1957). Population studies – Animal ecology and demography – Concluding remarks. Cold Spring Harbor Symposia on Quantitative Biology, 22, 415–427.

Iannarilli, F., Arnold, T. W., Erb, J., & Fieberg, J. R. (2019). Using lorelograms to measure and model correlation in binary data: Applications to ecological studies. Methods in Ecology and Evolution, 10, 2153–2162.

Klugkist, I., & Hoijtink, H. 2007. The Bayes factor for inequality and about equality constrained models. Computational Statistics & Data Analysis, 51, 6367–6379.

Kronfeld-Schor, N., & Dayan, T. (2003). Partitioning of time as an ecological resource. Annual review of ecology, evolution, and systematics, 34, 153–181.

Kronfeld-Schor, N., & Dayan, T. (2008). Activity patterns of rodents: the physiological ecology of biological rhythms. Biological Rhythm Research, 39, 193–211.

Levy, O., Dayan, T., Porter, W. P., & Kronfeld-Schor, N. (2019). Time and ecological resilience: can diurnal animals compensate for climate change by shifting to nocturnal activity?. Ecological Monographs, 89, e01334.

Magle, S. B., Fidino, M., Lehrer, E. W., Gallo, T., Mulligan, M. P., Ŕıos, M. J., … & Drake, D. (2019). Advancing urban wildlife research through a multi-city collaboration. Frontiers in Ecology and the Environment, 17, 232–239.

Michelot, T., & Blackwell, P. G. 2019. State-switching continuous-time correlated random walks. Methods in Ecology and Evolution, 10, 637–649.

Mittermeier, R. A., and Wilson, D. E. (2009). Handbook of the mammals of the world: vol. 1: carnivores. Lynx.

Moll, R. J., Cepek, J. D., Lorch, P. D., Dennis, P. M., Robison, T., Millspaugh, J. J., & Montgomery, R. A. (2018). Humans and urban development mediate the sympatry of competing carnivores. Urban Ecosystems, 21, 765–778.

Northrup, J. M., Vander Wal, E., Bonar, M., Fieberg, J., Laforge, M. P., Leclerc, M., Prokopenko, C. M., & Gerber, B. D. (2022). Conceptual and methodological advances in habitat-selection modeling: guidelines for ecology and evolution. Ecological Applications, 32, e02470.

Nowak, R. M., and Walker, E. P. (1999). Walker’s Mammals of the World (Vol. 1). Johns Hopkins University Press.

Pianka, E. R. (1973). The structure of lizard communities. Annual review of ecology and systematics, 4, 53–74.

R Core Team. (1923). R: A Language and Environment for Statistical Computing. R Foundation for Statistical Computing. Vienna, Austria. https://www.R-project.org/

Ridout, M. S., & Linkie, M. (2009). Estimating overlap of daily activity patterns from camera trap data. Journal of Agricultural, Biological, and Environmental Statistics, 14, 322–337.

Riede, S. J., van der Vinne, V., and Hut, R. A. (2017). The flexible clock: predictive and reactive homeostasis, energy balance and the circadian regulation of sleep–wake timing. Journal of Experimental Biology, 220, 738–749.

Rivera, K., Fidino, M., Farris, Z. J., Magle, S. B., Murphy, A., & Gerber, B. D. (2022). Rethinking habitat occupancy modeling and the role of diel activity in an anthropogenic world. The American Naturalist, 200(4), 556–570.

Schoener, T. W. (1974). Resource Partitioning in Ecological Communities: Research on how similar species divide resources helps reveal the natural regulation of species diversity. Science, 185(4145), 27–39.

Schoener, T. W. (1974). The compression hypothesis and temporal resource partitioning. Proceedings of the National Academy of Sciences, 71, 4169–4172.

Silvapulle, M. J., & Sen, P. K. (2011). Constrained statistical inference: Order, inequality, and shape constraints. John Wiley & Sons.

Sunarto, S., Kelly, M. J., Parakkasi, K., & Hutajulu, M. B. (2015). Cat coexistence in central Sumatra: ecological characteristics, spatial and temporal overlap, and implications for management. Journal of Zoology, 296, 104–115.

Tattersall, I. (2008). Avoiding commitment: cathemerality among primates. Biological Rhythm Research, 39, 213–228.

Thieurmel B., Elmarhraoui A. (2019) Suncalc: Compute sun position, sunlight phases, moon position and lunar phase. Available at: https://CRAN.R-project.org/package=suncalc.

Tucker, M. A., Böhning-Gaese, K., Fagan, W. F., Fryxell, J. M., Van Moorter, B., Alberts, S. C., … & Mueller, T. (2018). Moving in the Anthropocene: Global reductions in terrestrial mammalian movements. Science, 359, 466–469.

van der Veen, D. R., Riede, S. J., Heideman, P. D., Hau, M., van der Vinne, V., & Hut, R. A. (2017). Flexible clock systems: adjusting the temporal programme. Philosophical Transactions of the Royal Society B: Biological Sciences, 372, 20160254.

