## Supplemental Material 1 for "A model-based hypothesis framework to define and estimate the diel niche via the ‘Diel.Niche’ R package"

Running Head: Modeling diel hypotheses

**Key-words**: cathemeral, crepuscular, diel, Diel.Niche, diurnal, nocturnal, temporal niche

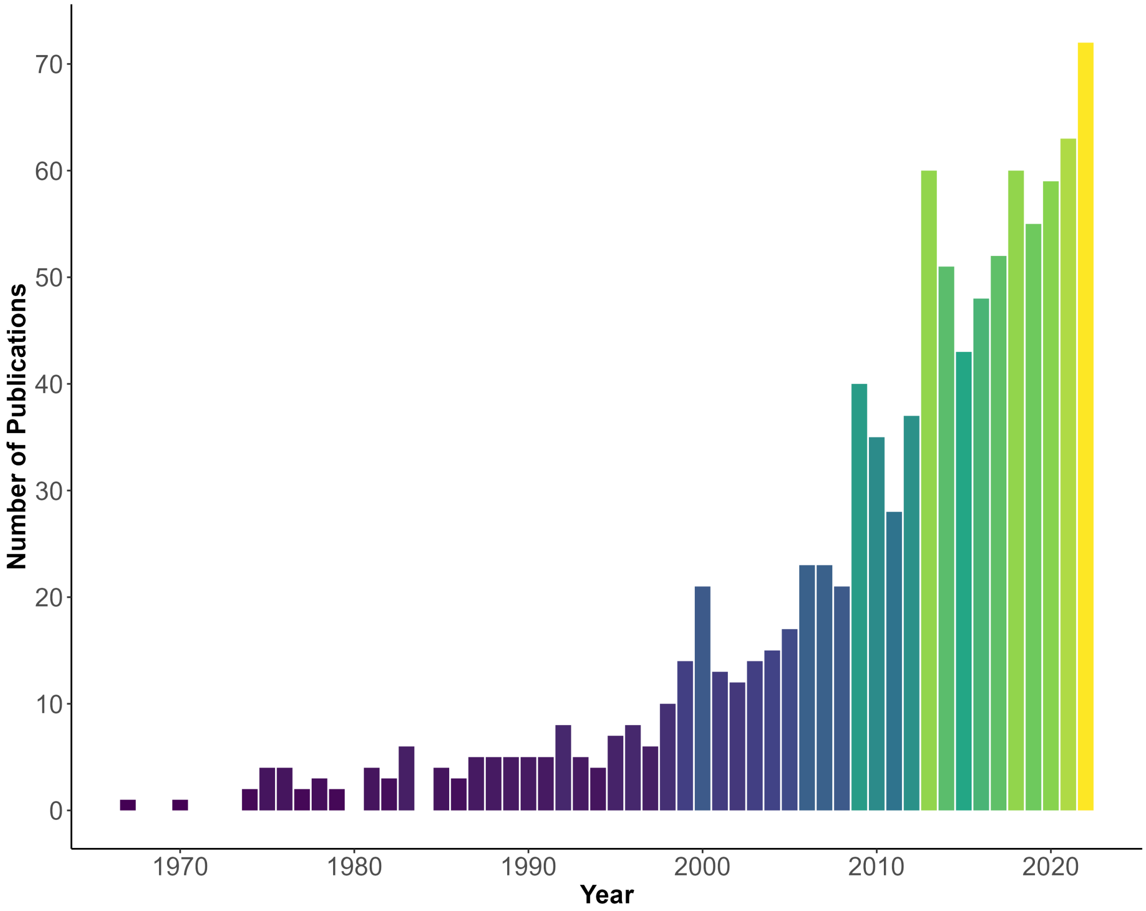


Figure S1: The number of publications per year from Scopus (https://www.scopus.com) using the search terms 'diel AND activity OR patterns AND animal'.

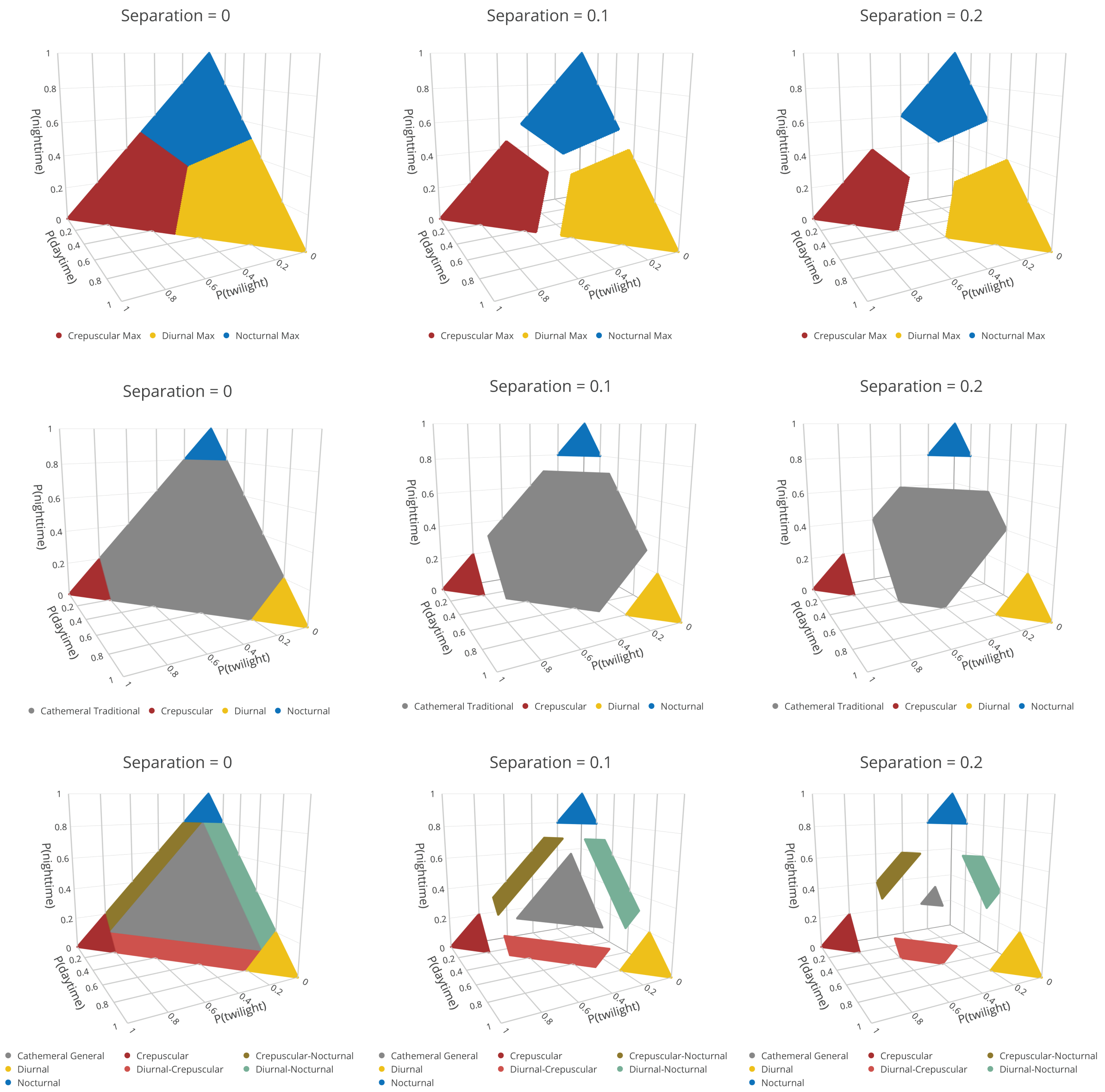


Figure S2: Hypotheses for different diel phenotypes with reduced parameter space for the simulation study. The parameter space was reduced to determine how model comparison was affected when the probabilities to simulate data could or could not be adjacent to other diel phenotypes.

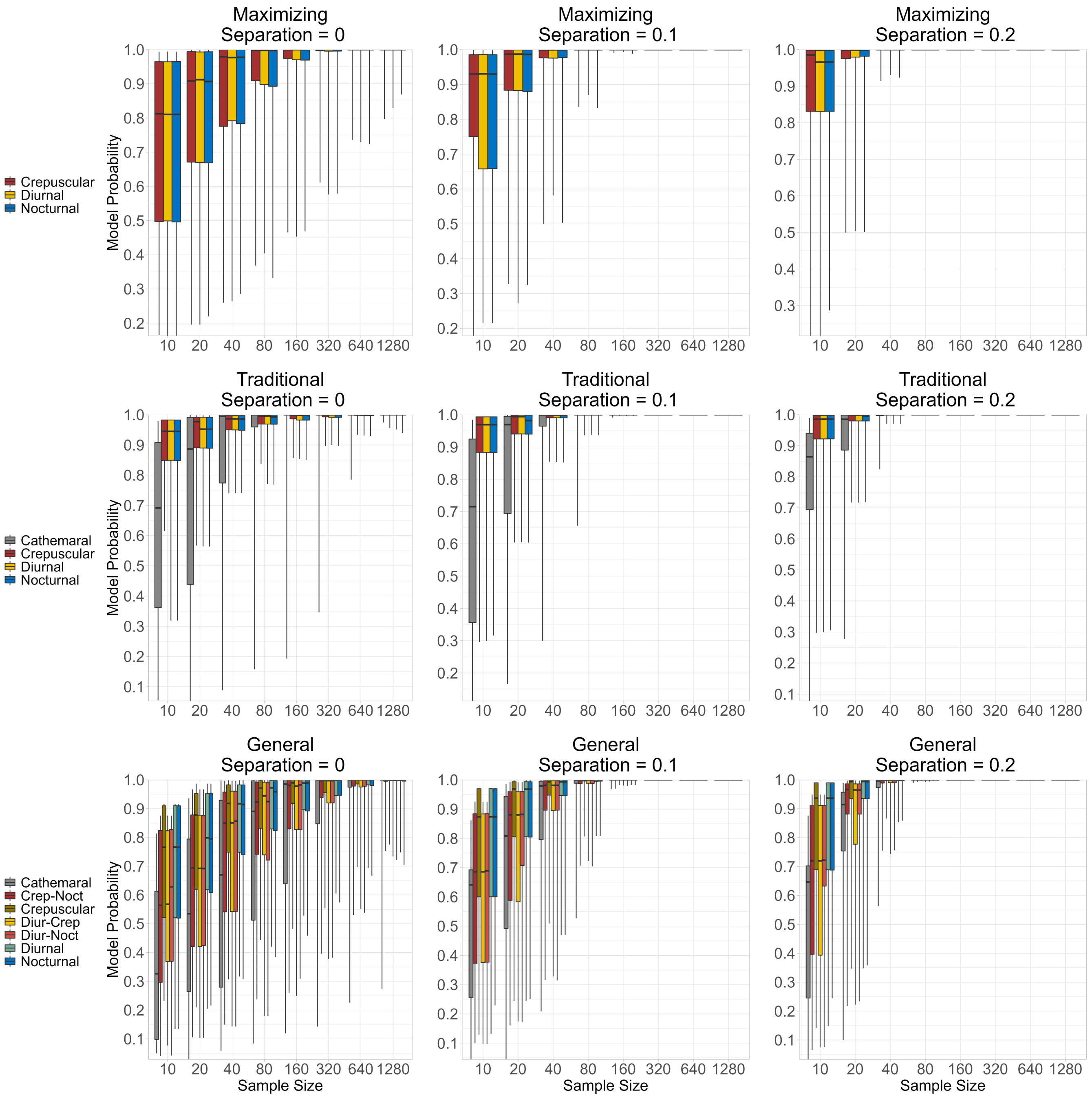


Figure S3: Simulation results for the Maximizing, Traditional, and General hypothesis sets. Each bar indicates the probability of support for the generating model across varying sample sizes and how much separation in probability space there is between hypotheses within a hypothesis set.

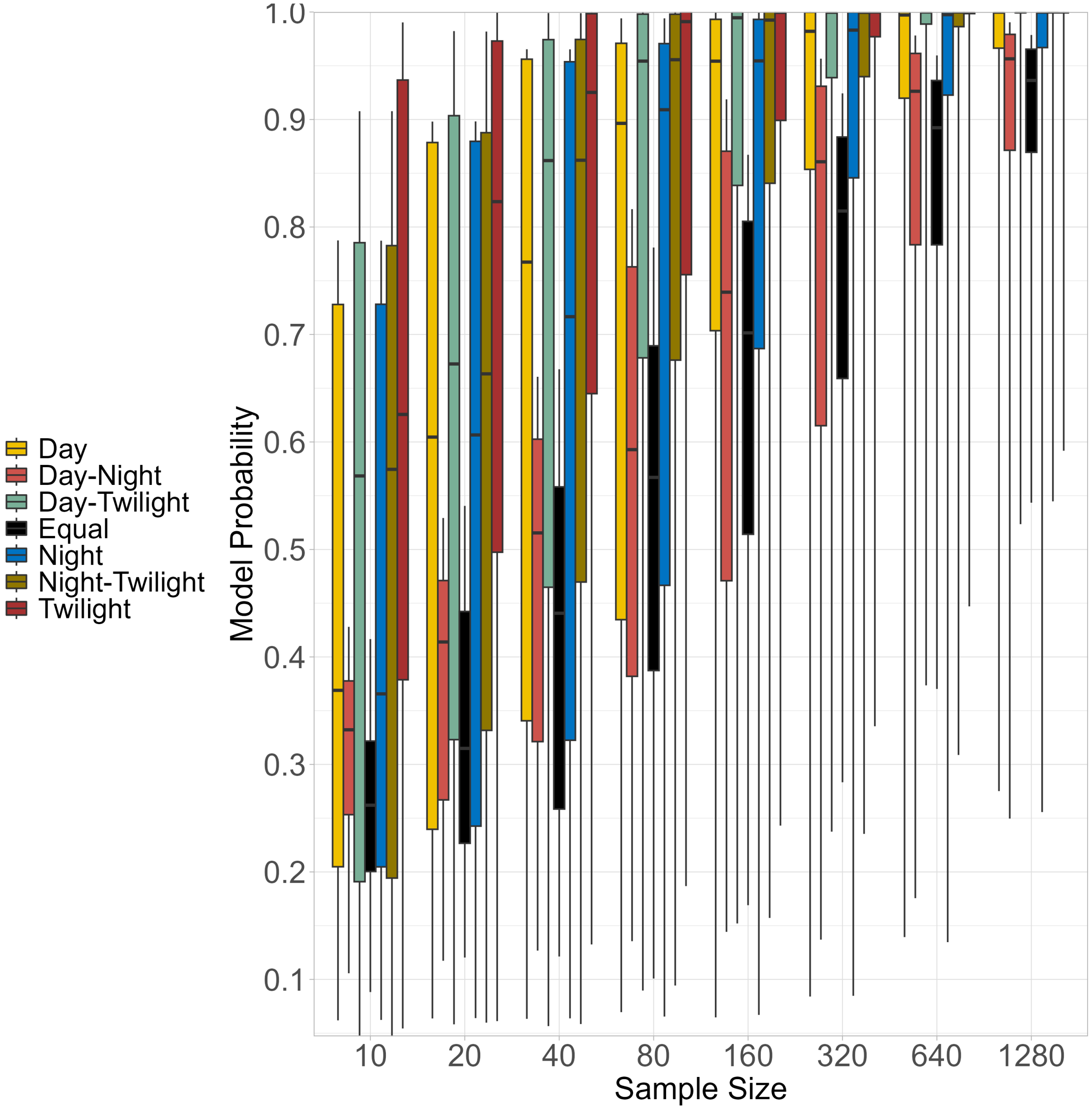


Figure S4: Simulation results for the Selection hypothesis set of the probability of support for the generating model across varying sample sizes.

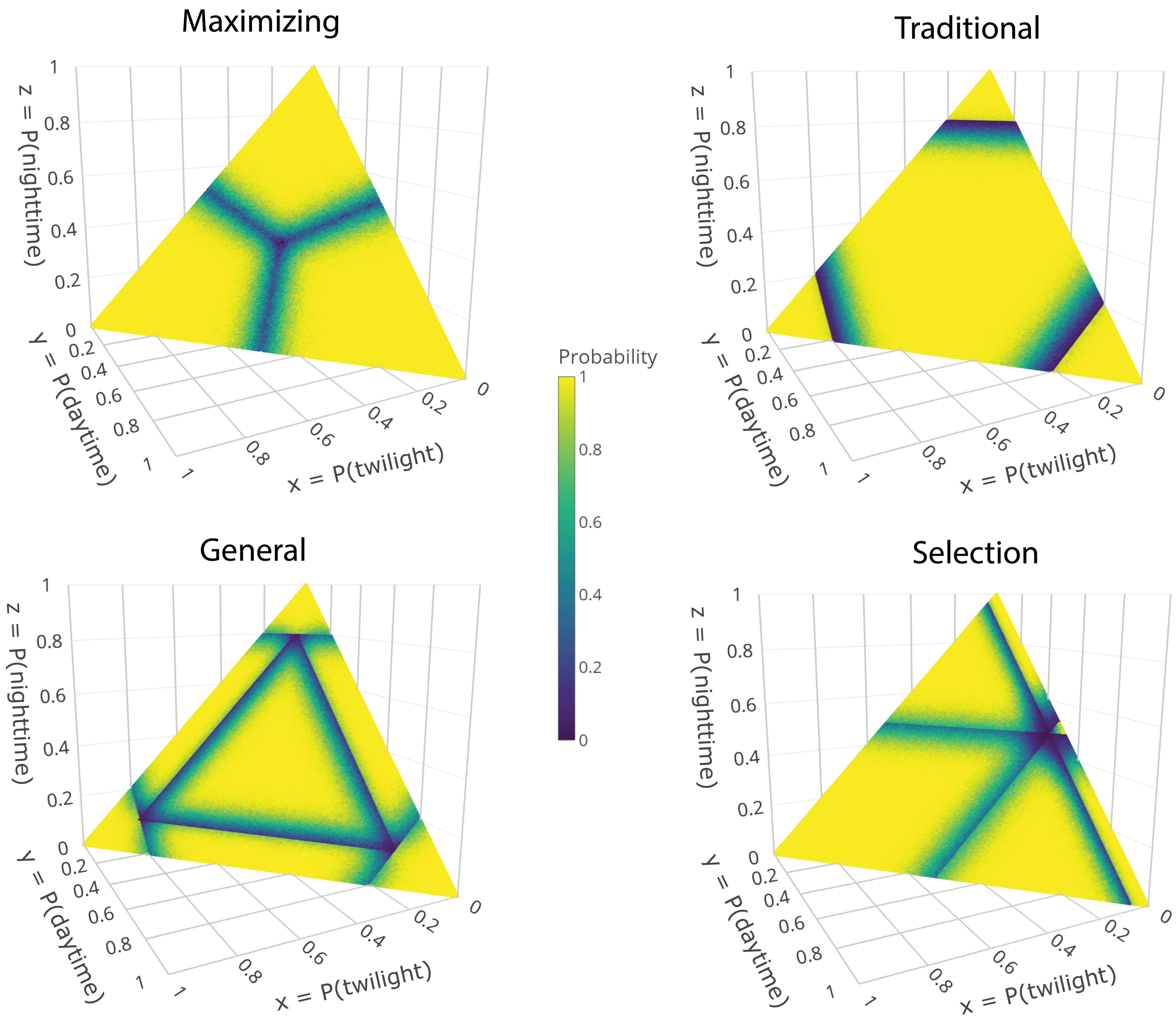


Figure S5: Simulation results depicting the probability that the generating model is most supported by parameter combinations (increments of 0.05) when the sample size is 100 total observations. These results indicate that it is more difficult to identify the data generating hypothesis when probabilities are near the bounds of other diel phenotypes.
