## Supplemental Material 2 for "A model-based hypothesis framework to define and estimate the diel niche via the ‘Diel.Niche’ R package"

#### Defining diel niche phenotypes

We define diel phenotype hypotheses using inequality and equality constraints on multinomial probabilities for estimation and model comparison in the R package `Diel.Niche`. This package provides wrapper functions to make use of the R package `multinomineq` (Heck and Davis-Stober, 2019). The hypotheses defined here are preset in the `Diel.Niche` package. The main phenotypes are defined by inequality constraints, but a few additional ones are specified as equalities.

Each hypothesis is defined by vector inequalities and equalities that constrain a vector of model probabilities $\boldsymbol{\theta}$. We assume the data are setup with ordered frequencies of detections in the twilight (dawn and dusk combined), daytime, and nighttime periods, such that there are two free model probabilities that can be constrained, as $\boldsymbol{\theta} = [p_{\text{tw}}, p_{\text{d}}]$ (the first two specified probabilities), while $p_{\text{n}}$ is derived as $1 - p_{\text{tw}} - p_{\text{d}}$.

Inequality constraints are defined by matrix $\mathbf{A}$ and vector $\mathbf{b}$, such that $\boldsymbol{A\theta} \leq \boldsymbol{b}$. Equality constraints are defined by matrix $\mathbf{C}$ and vector $\mathbf{d}$, such that $\boldsymbol{C\theta} = \boldsymbol{d}$, which is solved by sequential approximation of $|\boldsymbol{C\theta} - \boldsymbol{d}| < \delta$. Below, we specify parameters for the different models, including threshold ($\xi_1$, $\xi_2$), most probable values ($\eta$) and measures of variability ($\epsilon$). These parameters are preset in `Diel.Niche`, but can be changed easily. Below, where we specify an inequality that is less than a specific threshold value (e.g., $\xi_1$) and not less than or equal to that value, we include a very small amount of deviation ($\gamma$) so that we can still use the less than or equal framework. For example, $p_{\text{d}} < \xi_1$ is fit into this framework as

$p_d \leq \xi_1 - \gamma$, where $\gamma$ is a very small number (e.g., 0.0001).

Sections 1 to 5 outline complete hypotheses sets with either 3, 4, or 7 hypotheses. Sections 1-4 outline hypotheses sets (Maximizing, Traditional, General, and Selection) that define the complete parameter space (i.e., between 0 and 1 for each probability). Sections 5-6 are alternative hypotheses sets with more limited defined parameter space (Threshold, Variation). Section 7 includes descriptions for potentially useful extra hypotheses.

For each hypotheses below, we define it generally (*Hypothesis*), describe the inequalities (*Description*), and then define how the matrices and vectors need to be setup for implementing (*Implementation*). Note that the implementation is only ever in regards to $p_{tw}$ and $p_d$, as they are the two estimable parameters, whereas $p_n$ is derived. Therefore constraints on $p_n$ are in terms of the other two parameters. In the first subsection of sections 1-6, we write out a complete walkthrough of setting up, interpreting, and justifying the inequality constraints. The same math and logic is used throughout for all sections, but is not demonstrated after the first subsection.

### 1 Maximizing Hypotheses

The maximizing hypotheses set includes three hypotheses: diurnal, nocturnal, and crepuscular. This hypothesis set is setup to evaluate which time period is used the most. Since we are examining the most amount of activity in one time period there is no hypothesis about activity across multiple time periods (i.e., cathemeral).

Researchers may want to use this hypothesis set if they they are interested in determining the dominant diel period a species uses. As such, this hypothesis set only allows you to estimate unimodal patterns of diel activity.

#### 1.1 Diurnal

*Hypothesis*: An animal is primarily diurnal, such that the majority of its activity occurs during the daytime. It is an broad hypothesis without specifics about activity during the nighttime or twilight periods.

*Description*: The probability of activity in the daytime is greater than the probability of activity during twilight and nighttime. There are no constraints on how much activity during the daytime or any order between the probability of activity during the nighttime and twilight periods.

*Mathematical Inequalities*:

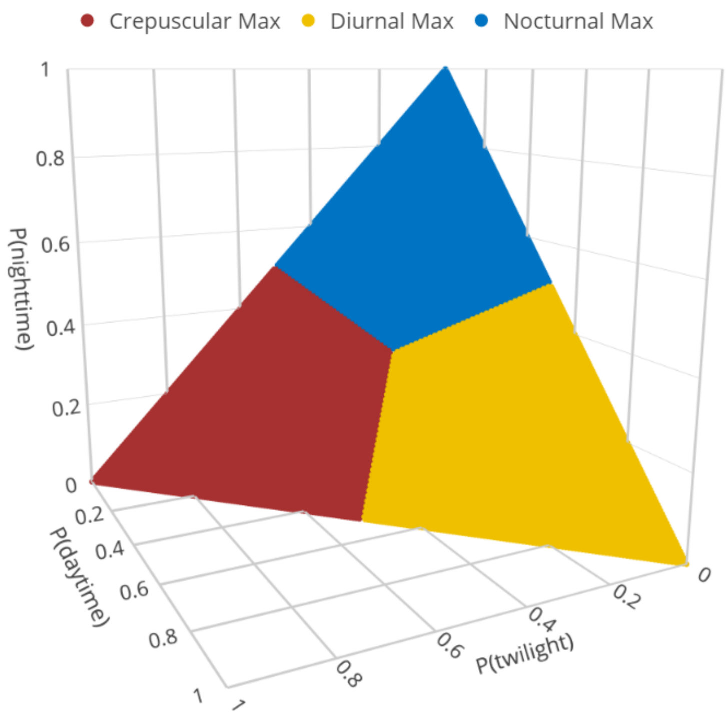


Figure 1: The defined parameter space for the three maximizing hypotheses.

$$\begin{aligned} p_{\text{tw}} &\leq p_{\text{d}} \\ p_{\text{n}} &\leq p_{\text{d}} \end{aligned} \tag{1}$$

*Implementation*: $\boldsymbol{A} = \begin{bmatrix} 1 & -1 \\ -1 & -2 \end{bmatrix}, \boldsymbol{b} = \begin{bmatrix} 0 \\ -1 \end{bmatrix}$

Therefore, all together we have,

$$\begin{aligned} \boldsymbol{A\theta} &\leq \boldsymbol{b} \\ \begin{bmatrix} (1) \times p_{\text{tw}} + (-1) \times p_{\text{d}} \\ (-1) \times p_{\text{tw}} + (-2) \times p_{\text{d}} \end{bmatrix} &\leq \begin{bmatrix} 0 \\ -1 \end{bmatrix} \\ \begin{bmatrix} p_{\text{tw}} - p_{\text{d}} \\ -p_{\text{tw}} - 2p_{\text{d}} \end{bmatrix} &\leq \begin{bmatrix} 0 \\ -1 \end{bmatrix} \end{aligned} \tag{2}$$

**Step-by-step**:
Interpretation Constraint 1:

$$\begin{aligned} & p_{\text{tw}} \leq p_{\text{d}} \\ & p_{\text{tw}} - p_{\text{d}} \leq 0, \text{ such that} \\ & (1) \times p_{\text{tw}} + (-1) \times p_{\text{d}} \leq 0 \end{aligned} \tag{3}$$

Interpretation Constraint 2:

$$\begin{aligned}
&p_{\text{n}} \leq p_{\text{d}} \\
&1 - p_{\text{tw}} - p_{\text{d}} \leq p_{\text{d}}, \text{ (given that } p_{\text{n}} = 1 - p_{\text{tw}} - p_{\text{d}}) \\
&1 - p_{\text{tw}} - p_{\text{d}} - p_{\text{d}} \leq 0 \\
&(-1) \times p_{\text{tw}} + (-2) \times p_{\text{d}} \leq -1
\end{aligned} \tag{4}$$

#### 1.2 Nocturnal

*Hypothesis*: An animal is primarily nocturnal, such that the majority of its activity occurs during the nighttime. It is a broad hypothesis without specifics about activity during the daytime or twilight periods.

*Description*: The probability of activity in the nighttime is greater than the probability of activity during the daytime and twilight. There are no constraints on how much activity during the nighttime or any order between the probability of activity during the daytime and twilight periods.

*Mathematical Inequalities*:

$$\begin{aligned}
p_{\text{d}} &\leq p_{\text{n}} \\
p_{\text{tw}} &\leq p_{\text{n}}
\end{aligned} \tag{5}$$

*Implementation*: $\boldsymbol{A} = \begin{bmatrix} 1 & 2 \\ 2 & 1 \end{bmatrix}, \boldsymbol{b} = \begin{bmatrix} 1 \\ 1 \end{bmatrix}$

#### 1.3 Crepuscular

*Hypothesis*: An animal is primarily crepuscular, such that the majority of its activity occurs during the twilight. It is a broad hypothesis without specifics about activity during the daytime or nighttime periods.

*Description*: The probability of activity in the twilight period is greater than the probability of activity during the daytime or nighttime. There are no constraints on the amount of activity during twilight or any order between the probability of activity during the daytime and nighttime periods.

*Mathematical Inequalities*:

$$\begin{aligned}
p_{\text{d}} &\leq p_{\text{tw}} \\
p_{\text{n}} &\leq p_{\text{tw}}
\end{aligned} \tag{6}$$

*Implementation*: $\boldsymbol{A} = \begin{bmatrix} -1 & 1 \\ -2 & -1 \end{bmatrix}, \boldsymbol{b} = \begin{bmatrix} 0 \\ -1 \end{bmatrix}$

### 2 Traditional Hypotheses

The traditional hypotheses set includes four hypotheses: diurnal, nocturnal, crepuscular, and cathemeral. Diurnal, nocturnal, and crepuscular are each defined by having a large amount of activity in each of their respective periods of time that is more than a defined lower threshold ($\xi_1$). If an animal is not mostly active in one time period it is then defined as cathemeral. This occurs when either two or three time periods are used more than $1-\xi_1$. Note that in `Diel.Niche` $\xi_1$ is the first element of the variable 'xi'.

Researchers may want to use this hypothesis set if they want to evaluate the fit of standard diel phenotypes in the literature and provide uncertainty estimates to a specific diel phenotype when reporting a species' activity pattern in a research study.

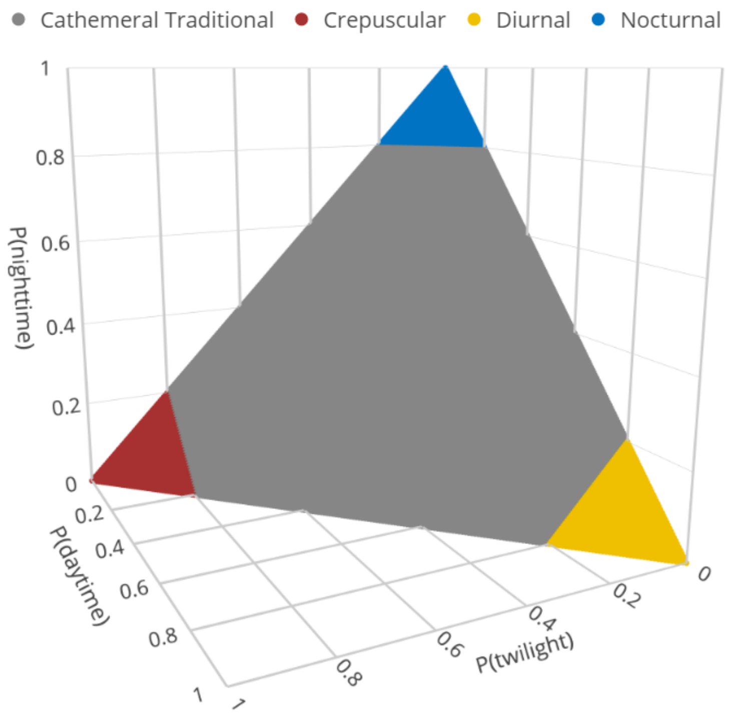


Figure 2: Defined parameter space for the four traditional hypotheses. The parameter $\xi_1$ is set to 0.8, indicating that to be diurnal, crepuscular, or nocturnal the animal needs to have a probability of use in each respective time period of 0.8 or greater. If an animal has less than this amount in daytime, nighttime, and twilight it is considered cathemeral.

#### 2.1 Diurnal

*Hypothesis*: An animal is primarily diurnal, such that most of its activity occurs during the daytime and above a lower probability threshold ($\xi_1$). If there is additional activity during the night and twilight, it is a combined amount less than 1-$\xi_1$.

*Description*: The probability of activity in the daytime is greater than or equal to the probability $\xi_1$.

*Mathematical Inequalities*:

$$p_{\mathrm{d}} \geq \xi_1 \tag{7}$$

*Implementation*: $\boldsymbol{A} = \begin{bmatrix} 0 & -1 \end{bmatrix}, \boldsymbol{b} = \begin{bmatrix} -\xi_1 \end{bmatrix}$

Therefore, all together we have,

$$\begin{aligned} \boldsymbol{A\theta} &\leq \boldsymbol{b} \\ \left[(0) \times p_{\mathrm{tw}} + (-1) \times p_{\mathrm{d}}\right] &\leq \left[-\xi_1\right] \\ \left[-p_{\mathrm{d}}\right] &\leq \left[-\xi_1\right] \\ \left[p_{\mathrm{d}}\right] &\geq \left[\xi_1\right] \end{aligned} \tag{8}$$

**Step-by-step**:
Interpretation Constraint 1:

$$\begin{aligned} & p_d \geq \xi_1 \\ & -p_d \leq -\xi_1, \text{ such that} \\ & (0) \times p_{\mathrm{tw}} + (-1) \times p_{\mathrm{d}} \leq -\xi_1 \end{aligned} \tag{9}$$

#### 2.2 Nocturnal

*Hypothesis*: An animal is primarily nocturnal, such that most of its activity occurs during the nighttime and above a probability threshold ($\xi_1$). If there is additional activity during daytime and twilight, it is a combined amount of less than 1-$\xi_1$.

*Description*: The probability of activity in the nighttime is great than or equal to the probability threshold $\xi_1$.

*Mathematical Inequalities*:

$$p_{\mathrm{n}} \geq \xi_1 \tag{10}$$

*Implementation*: $\boldsymbol{A} = \begin{bmatrix} 1 & 1 \end{bmatrix}, \boldsymbol{b} = \begin{bmatrix} -\xi_1 + 1 \end{bmatrix}$

#### 2.3 Crepuscular

*Hypothesis*: An animal is primarily crepuscular, such that most of its activity occurs during twilight and above a lower probability threshold of $\xi_1$. If there is additional activity during the nighttime and daytime, it is a combined amount of less than 1-$\xi_1$.

*Description*: The probability of activity in the twilight is great than or equal to the threshold $\xi_1$.

*Mathematical Inequalities*:

$$p_{\mathrm{tw}} \geq \xi_1 \tag{11}$$

*Implementation*: $\boldsymbol{A} = \begin{bmatrix} -1 & 0 \end{bmatrix}, \boldsymbol{b} = \begin{bmatrix} -\xi_1 \end{bmatrix}$

#### 2.4 Cathemeral

*Hypothesis*: An animal uses two or three diel periods (daytime, nighttime, twilight) and never uses one period more than the probability threshold of $\xi_1$.

*Description*: The probability of activity in the daytime, nighttime, and twilight periods is less than $\xi_1$.

*Mathematical Inequalities*:

$$\begin{aligned} p_{\mathrm{d}} &< \xi_1 \\ p_{\mathrm{tw}} &< \xi_1 \\ p_{\mathrm{n}} &< \xi_1 \end{aligned} \tag{12}$$

*Implementation*: $\boldsymbol{A} = \begin{bmatrix} 0 & 1 \\ 1 & 0 \\ -1 & -1 \end{bmatrix}, \boldsymbol{b} = \begin{bmatrix} \xi_1 - \gamma \\ \xi_1 - \gamma \\ \xi_1 - \gamma - 1 \end{bmatrix}$

Note that $\gamma$ is a pre-specified very small number. It is subtracted from $\xi_1$ because we are using a less than or equal to mathematical setup and we need this inequality to specify the probabilities to be less than $\xi_1$.

### 3 General Hypotheses

The general hypothesis set includes three of the traditional hypotheses (diurnal, nocturnal, and crepuscular) and are not repeated in this section. However, this hypothesis set modifies the cathemeral hypothesis to be more specific. To do so, we need to consider a probability threshold where a time period cannot be used more than $\xi_1$ and is not used less than a lower probability threshold $\xi_2$. Note that in `Diel.Niche` $\xi_1$ and $\xi_2$ are specified as the first and second elements of the variable 'xi', respectively.

Researchers may want to use this hypothesis set if they want to determine bimodal activity patterns a species may have. For example, this is the only hypothesis set that can evaluate if a species uses both twilight and day more than some set threshold (i.e., Diurnal-Crepuscular). As the hypothesis set with the most competing hypotheses, researchers should have a substantial sample size when using it (see manuscript for suggestions).

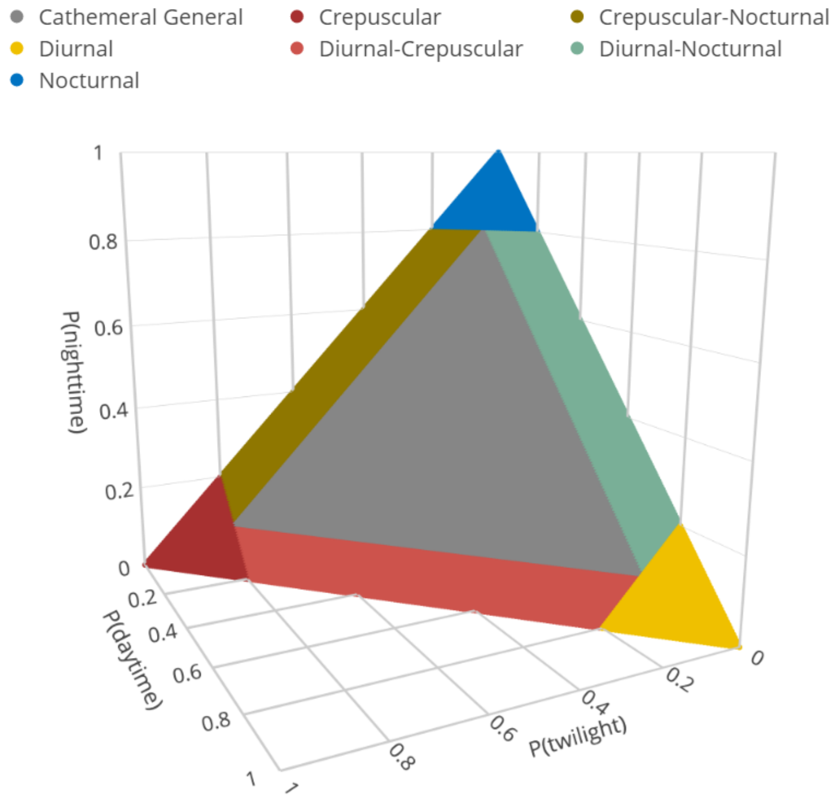


Figure 3: The defined parameter space for the seven General hypotheses. The parameter $\xi_1$ is set to 0.8, indicating that to be diurnal, crepuscular, or nocturnal the animal needs to have a probability of use in each respective time period of 0.8 or greater. The parameter $\xi_2$ is set to 0.1, indicating that an animal is cathemeral when all three diel periods are used more than this amount. When only two diel periods are used more than this amount then it is either Crepuscular-Nocturnal, Diurnal-Crepuscular, or Diurnal-Nocturnal.

##### 3.1 Cathemeral

*Hypothesis*: An animal uses all three diel periods, but not more than with probability $\xi_1$ and no less than probability $\xi_2$.

*Description*: The probability of daytime, twilight, and nocturnal activity is less than $\xi_1$ and more than $\xi_2$.

*Mathematical Inequalities*:

$$\begin{aligned} p_{\mathrm{d}} &\geq \xi_2 \\ p_{\mathrm{n}} &\geq \xi_2 \\ p_{\mathrm{tw}} &\geq \xi_2 \\ p_{\mathrm{d}} &< \xi_1 \\ p_{\mathrm{n}} &< \xi_1 \end{aligned} \tag{13}$$

*Implementation*: $\boldsymbol{A} = \begin{bmatrix} 0 & -1 \\ 1 & 1 \\ -1 & 0 \\ 0 & 1 \\ -1 & -1 \end{bmatrix}, \boldsymbol{b} = \begin{bmatrix} -\xi_1 \\ -\xi_2 + 1 \\ -\xi_2 \\ \xi_1 - \gamma \\ \xi_1 - \gamma - 1 \end{bmatrix}$

Therefore, all together we have,

$$\begin{gathered} \boldsymbol{A\theta} \leq \boldsymbol{b} \\ \begin{bmatrix} (0) \times p_{\mathrm{tw}} + (-1) \times p_{\mathrm{d}} \\ (1) \times p_{\mathrm{tw}} + (1) \times p_{\mathrm{d}} \\ (-1) \times p_{\mathrm{tw}} + (0) \times p_{\mathrm{d}} \\ (0) \times p_{\mathrm{tw}} + (1) \times p_{\mathrm{d}} \\ (-1) \times p_{\mathrm{tw}} + (-1) \times p_{\mathrm{d}} \end{bmatrix} \leq \begin{bmatrix} -\xi_2 \\ -\xi_2 + 1 \\ -\xi_2 \\ \xi_1 - \gamma \\ \xi_1 - \gamma + 1 \end{bmatrix} \\ \begin{bmatrix} -p_{\mathrm{d}} \\ p_{\mathrm{tw}} + p_{\mathrm{d}} \\ -p_{\mathrm{tw}} \\ p_{\mathrm{d}} \\ -p_{\mathrm{tw}} - p_{\mathrm{d}} \end{bmatrix} \leq \begin{bmatrix} -\xi_2 \\ -\xi_2 + 1 \\ -\xi_2 \\ \xi_1 - \gamma \\ \xi_1 - \gamma - 1 \end{bmatrix} \end{gathered} \tag{14}$$

**Step-by-step**:

Interpretation Constraint 1:

$$\begin{aligned} & p_d \geq \xi_2 \\ & -p_d \leq -\xi_2, \text{ such that} \\ & (0) \times p_{\mathrm{tw}} + (-1) \times p_{\mathrm{d}} \leq -\xi_2 \end{aligned} \tag{15}$$

Interpretation Constraint 2:

$$\begin{aligned}
&p_n > \xi_2 \\
&1 - p_{\text{tw}} - p_d > \xi_2, \text{ where } p_{\text{n}} = 1 - p_{\text{tw}} - p_{\text{d}} \\
&- p_{\text{tw}} - p_d > \xi_2 - 1, \\
&p_{\text{tw}} + p_d < -\xi_2 + 1, \text{ such that} \\
&(1) \times p_{\text{tw}} + (1) \times p_{\text{d}} < -\xi_2 + 1
\end{aligned} \tag{16}$$

Interpretation Constraint 3:

$$\begin{aligned}
&p_{\text{tw}} > \xi_2 \\
&- p_{\text{tw}} < -\xi_2, \text{ such that} \\
&(-1) \times p_{\text{tw}} + (0) \times p_{\text{d}} < -\xi_2
\end{aligned} \tag{17}$$

Interpretation Constraint 4:

$$\begin{aligned}
&p_d < \xi_1, \text{ where} \\
&p_d \leq \xi_1 - \gamma, \text{ such that} \\
&(0) \times p_{\text{tw}} + (1) \times p_{\text{d}} < \xi_1 - \gamma
\end{aligned} \tag{18}$$

Interpretation Constraint 5:

$$\begin{aligned}
&p_n < \xi_1, \text{where} \\
&p_n \leq \xi_1 - \gamma, \text{ and } p_{\text{n}} = 1 - p_{\text{tw}} - p_{\text{d}} \\
&1 - p_{\text{tw}} - p_{\text{d}} \leq \xi_1 - \gamma \\
&- p_{\text{tw}} - p_{\text{d}} \leq \xi_1 - \gamma - 1, \text{ such that} \\
&(-1) \times p_{\text{tw}} + (-1) \times p_{\text{d}} \leq \xi_1 - \gamma - 1
\end{aligned} \tag{19}$$

#### 3.2 Diurnal-Crepuscular

*Hypothesis*: An animal uses both twilight and daytime with probabilities more than $\xi_2$ and less than $\xi_1$. Nighttime activity is used very little with less than probability $\xi_2$.

*Description*: The probability of nighttime activity is lower than $\xi_2$, while the probability of activity in twilight and daytime is above $\xi_2$ and less than $\xi_1$.

*Mathematical Inequalities*:

$$\begin{aligned}
&p_{\text{n}} \leq \xi_2 \\
&p_{\text{tw}} \geq \xi_2 \\
&p_{\text{tw}} \leq \xi_1 \\
&p_{\text{d}} \geq \xi_2 \\
&p_{\text{d}} \leq \xi_1
\end{aligned} \tag{20}$$

*Implementation*: $\boldsymbol{A} = \begin{bmatrix} -1 & -1 \\ -1 & 0 \\ 1 & 0 \\ 0 & -1 \\ 0 & 1 \end{bmatrix}, \boldsymbol{b} = \begin{bmatrix} \xi_2 - 1 \\ -\xi_2 \\ \xi_1 \\ -\xi_2 \\ \xi_1 \end{bmatrix}$

#### 3.3 Diurnal-Nocturnal

*Hypothesis*: An animal uses both nighttime and daytime with probabilities more than $\xi_2$ and less than $\xi_1$. Twilight activity is used very little with less than probability $\xi_2$.

*Description*: The probability of twilight activity is lower than $\xi_2$, while the probability of activity in nighttime and daytime is above $\xi_2$ and less than $\xi_1$.

*Mathematical Inequalities*:

$$\begin{aligned} p_{\mathrm{tw}} &\leq \xi_2 \\ p_d &\geq \xi_2 \\ p_d &\leq \xi_1 \\ p_{\mathrm{n}} &\geq \xi_2 \\ p_{\mathrm{n}} &\leq \xi_1 \end{aligned} \tag{21}$$

*Implementation*: $\boldsymbol{A} = \begin{bmatrix} 1 & 0 \\ 0 & -1 \\ 0 & 1 \\ 1 & 1 \\ -1 & -1 \end{bmatrix}, \boldsymbol{b} = \begin{bmatrix} \xi_2 \\ -\xi_2 \\ \xi_1 \\ -\xi_2 + 1 \\ \xi_1 - 1 \end{bmatrix}$

#### 3.4 Crepuscular-Nocturnal

*Hypothesis*: An animal uses both nighttime and twilight with probabilities more than $\xi_2$ and less than $\xi_1$. Daytime activity is used very little with a probability less than $\xi_2$

*Description*: The probability of daytime activity is lower than $\xi_2$, while the probability of activity in nighttime and twilight is above $\xi_2$ and less than $\xi_1$

*Mathematical Inequalities*:

$$\begin{aligned} p_d &\leq \xi_2 \\ p_n &\geq \xi_2 \\ p_n &\leq \xi_1 \\ p_{\mathrm{tw}} &\geq \xi_2 \\ p_{\mathrm{tw}} &\leq \xi_1 \end{aligned} \tag{22}$$

*Implementation*: $\boldsymbol{A} = \begin{bmatrix} 0 & 1 \\ 1 & 1 \\ -1 & -1 \\ -1 & 0 \\ - & 0 \end{bmatrix}, \boldsymbol{b} = \begin{bmatrix} \xi_2 \\ -\xi_2 + 1 \\ \xi_1 - 1 \\ -\xi_2 \\ \xi_1 \end{bmatrix}$

### 4 Selection Hypotheses

The selection hypotheses are day, night, twilight, day-night, night-twilight, day-twilight, and selection-available. With all other hypotheses we have focused on estimating the amount of activity use in each diel period and calling it a certain diel phenotype. Here, we focus on how an animal uses the periods, given the available amount of time in each period.

Researchers may want to use this hypothesis set if they are interested in certain diel periods being used more than they are available, i.e., 'selected'. This is particularly relevant to the use of the twilight period, which will generally have proportionally less time in this period than the day or night periods (excluding summer and winter times of the Arctic and Antarctic).

We require proportional availability by diel period to be supplied to evaluate these hypotheses. Let's define $\mathbf{p}_{\text{avail}} = [p_{\text{av.tw}},\ p_{\text{av.d}}]$. For example, $\mathbf{p}_{\text{avail}} = [0.04\ \ 0.48]$, thus implying that $p_{\text{av.n}} = 1 - 0.04 - 0.48 = 0.48$.

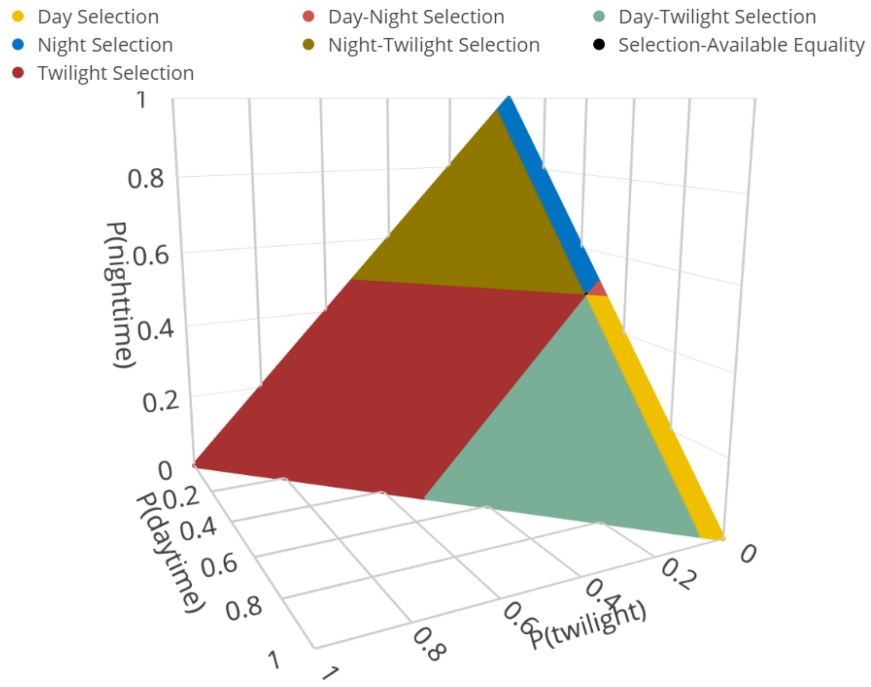


Figure 4: Defined parameter space for the seven selection hypotheses. The parameter space is defined where $\mathbf{p}_{\text{avail}} = [0.04\ \ 0.48]$.

#### 4.1 Day Selection

*Hypothesis*: An animal uses the day diel period more than available, and uses the night and twilight period in proportion or less than available.

*Description*: The proportion of activity use to available for the day period ($p_{\mathrm{d}}/p_{\mathrm{av.d}}$) is greater than one. Further, the proportion of activity use to available for the night and twilight are less than or equal to one.

*Mathematical Inequalities*:

$$\begin{aligned} &p_{\mathrm{d}}/p_{\mathrm{av.d}} > 1 \\ &p_{\mathrm{tw}}/p_{\mathrm{av.tw}} \leq 1 \\ &p_{\mathrm{n}}/p_{\mathrm{av.n}} \leq 1 \end{aligned} \tag{23}$$

*Implementation*: $\boldsymbol{A} = \begin{bmatrix} 0 & -1/p_{\mathrm{av.d}} \\ 1/p_{\mathrm{av.tw}} & 0 \\ -1 & -1 \end{bmatrix}, \boldsymbol{b} = \begin{bmatrix} -1-\gamma \\ 1 \\ p_n - 1 \end{bmatrix}$

Therefore, all together we have,

$$\begin{aligned} \boldsymbol{A\theta} &\leq \boldsymbol{b} \\ \begin{bmatrix} (0)\times p_{\mathrm{tw}} + (-1/p_{\mathrm{av.d}})\times p_{\mathrm{d}} \\ (1/p_{\mathrm{av.tw}})\times p_{\mathrm{tw}} + (0)\times p_{\mathrm{d}} \\ (-1)\times p_{\mathrm{tw}} + (-1)\times p_{\mathrm{d}} \end{bmatrix} &\leq \begin{bmatrix} -1-\gamma \\ 1 \\ p_{\mathrm{av.tw}} - p_{\mathrm{av.d}} \end{bmatrix} \\ \begin{bmatrix} (p_{\mathrm{tw}}/p_{\mathrm{av.tw}}) \\ (p_{\mathrm{d}}/p_{\mathrm{av.d}}) \\ -p_{\mathrm{tw}} - p_{\mathrm{d}} \end{bmatrix} &\leq \begin{bmatrix} -1-\gamma \\ 1 \\ -p_{\mathrm{av.tw}} - p_{\mathrm{av.d}} \end{bmatrix} \end{aligned} \tag{24}$$

**Step-by-step**:
Interpretation Constraint 1:

$$\begin{aligned} &p_d/p_{\mathrm{av.d}} > 1 \text{ where,} \\ &p_d/p_{\mathrm{av.d}} \geq 1+\gamma \\ &-p_d/p_{\mathrm{av.d}} \leq -1-\gamma \\ &(0)\times p_{\mathrm{tw}} + (-1/p_{\mathrm{av.d}})\times p_d \leq -1-\gamma \end{aligned} \tag{25}$$

Interpretation Constraint 2:

$$\begin{aligned} &p_{\mathrm{tw}}/p_{\mathrm{av.tw}} \leq 1 \\ &(1/p_{\mathrm{av.tw}})\times p_{\mathrm{tw}} + (0)\times p_d \leq 1 \end{aligned} \tag{26}$$

Interpretation Constraint 3:

$$\begin{aligned} &p_{\mathrm{n}}/p_{\mathrm{av.n}} \leq 1 \\ &(1 - p_{\mathrm{tw}} - p_{\mathrm{d}})/p_{\mathrm{av.n}} \leq 1 \\ &1 - p_{\mathrm{tw}} - p_{\mathrm{d}} \leq p_{\mathrm{av.n}} \\ &-p_{\mathrm{tw}} - p_{\mathrm{d}} \leq p_{\mathrm{av.n}} - 1 \\ &(-1)\times p_{\mathrm{tw}} + (-1)\times p_d \leq p_{\mathrm{av.n}} - 1 \end{aligned} \tag{27}$$

#### 4.2 Night Selection

*Hypothesis*: An animal uses the night diel period more than available, and uses the day and twilight period in proportion or less than available.

*Description*: The proportion of activity use to available for the night period, $p_{\text{n}}/p_{\text{av.n}}$, is greater than one. Further, the proportion of activity use to available for the day and twilight are less than or equal to one.

*Mathematical Inequalities*:

$$\begin{aligned} p_{\text{n}}/p_{\text{av.n}} &> 1 \\ p_{\text{tw}}/p_{\text{av.tw}} &\leq 1 \\ p_{\text{d}}/p_{\text{av.d}} &\leq 1 \end{aligned} \tag{28}$$

*Implementation*: $\boldsymbol{A} = \begin{bmatrix} 1 & 1 \\ 1/p_{\text{av.tw}} & 0 \\ 0 & 1/p_{\text{av.d}} \end{bmatrix}, \boldsymbol{b} = \begin{bmatrix} -1 - \gamma \times p_{\text{av.n}} + 1 \\ 1 \\ 1 \end{bmatrix}$

#### 4.3 Twilight Selection

*Hypothesis*: An animal uses the twilight diel period more than available, and uses the day and night period in proportion or less than available.

*Description*: The proportion of activity use to available for the twilight period, $p_{\text{tw}}/p_{\text{av.tw}}$, is greater than one. Further, the proportion of activity use to available for the day and night are less than or equal to one.

*Mathematical Inequalities*:

$$\begin{aligned} p_{\text{tw}}/p_{\text{av.tw}} &> 1 \\ p_{d}/p_{\text{av.d}} &\leq 1 \\ p_{\text{n}}/p_{\text{av.n}} &\leq 1 \end{aligned} \tag{29}$$

*Implementation*: $\boldsymbol{A} = \begin{bmatrix} -1/p_{\text{av.tw}} & 0 \\ 0 & 1/p_{\text{av.d}} \\ -1 & -1 \end{bmatrix}, \boldsymbol{b} = \begin{bmatrix} -1 - \gamma \\ 1 \\ p_{\text{av.n}} - 1 \end{bmatrix}$

#### 4.4 Day-Twilight Selection

*Hypothesis*: An animal uses both the day and twilight diel periods more than available, and uses the night period in proportion to or less than available.

*Description*: The proportion of activity use to available for the day period ($p_{\text{d}}/p_{\text{av.d}}$) and twilight period ($p_{\text{tw}}/p_{\text{av.tw}}$), is greater than one. Further, the proportion of activity use to available for the night is less than or equal to one.

*Mathematical Inequalities*:

$$\begin{aligned} &p_{\text{d}}/p_{\text{av.d}} > 1 \\ &p_{\text{tw}}/p_{\text{av.tw}} > 1 \\ &p_{\text{n}}/p_{\text{av.n}} \leq 1 \end{aligned} \tag{30}$$

*Implementation*: $\boldsymbol{A} = \begin{bmatrix} 0 & -1/p_{\text{av.d}} \\ -1/p_{\text{av.tw}} & 0 \\ -1 & -1 \end{bmatrix}, \boldsymbol{b} = \begin{bmatrix} -1-\gamma \\ -1-\gamma \\ p_{\text{av.n}}-1 \end{bmatrix}$

#### 4.5 Night-Twilight Selection

*Hypothesis*: An animal uses both the night and twilight diel periods more than available, and uses the day period in proportion or less than available.

*Description*: The proportion of activity use to available for the night period ($p_n/p_{\text{av.n}}$) and twilight period ($p_{\text{tw}}/p_{\text{av.tw}}$), is greater than one. Further, the proportion of activity use to available for the day is less than or equal to one.

*Mathematical Inequalities*:

$$\begin{aligned} &p_{\text{n}}/p_{\text{av.n}} > 1 \\ &p_{\text{tw}}/p_{\text{av.tw}} > 1 \\ &p_{\text{d}}/p_{\text{av.d}} \leq 1 \end{aligned} \tag{31}$$

*Implementation*: $\boldsymbol{A} = \begin{bmatrix} 1 & 1 \\ -1/p_{\text{av.tw}} & 0 \\ 0 & 1/p_{\text{av.d}} \end{bmatrix}, \boldsymbol{b} = \begin{bmatrix} -1-\gamma \times p_{\text{av.n}}+1 \\ -1+\gamma \\ 1 \end{bmatrix}$

#### 4.6 Day-Night Selection

*Hypothesis*: An animal uses both the day and night diel periods more than available, and uses the twilight period in proportion or less than available.

*Description*: The proportion of activity use to available for the day period ($p_d/p_{\text{av.d}}$) and night period ($p_n/p_{\text{av.n}}$), is greater than one. Further, the proportion of activity use to available for twilight is less than or equal to one.

*Mathematical Inequalities*:

$$\begin{aligned} &p_{\text{d}}/p_{\text{av.d}} > 1 \\ &p_{\text{n}}/p_{\text{av.n}} > 1 \\ &p_{\text{tw}}/p_{\text{av.tw}} \leq 1 \end{aligned} \tag{32}$$

*Implementation*: $\boldsymbol{A} = \begin{bmatrix} 0 & -1/p_{\text{av.d}} \\ 1 & 1 \\ 1/p\text{av.tw} & 0 \end{bmatrix}, \boldsymbol{b} = \begin{bmatrix} -1-\gamma \\ -1-\gamma \times p_{\text{av.n}} + 1 \\ 1 \end{bmatrix}$

#### 4.7 Use is in Proportion to Availability Hypothesis

*General Hypothesis*: An animal is active in proportion to the available time in each diel period.

*Specific Hypothesis*: Daytime, nighttime, and crepuscular activity are equal to their availability.

Mathematical Inequalities:

$$\begin{aligned} p_{\text{tw}} &= p_{\text{av.tw}} \\ p_{\text{d}} &= p_{\text{av.d}} \\ p_{\text{n}} &= p_{\text{av.n}} \end{aligned} \tag{33}$$

Implementation: $\boldsymbol{C} = \begin{bmatrix} 1 & 0 \\ 0 & 1 \end{bmatrix}, \boldsymbol{d} = \begin{bmatrix} p_{\text{av.tw}} \\ p_{\text{av.d}} \end{bmatrix}$

### 5 Threshold Hypotheses

The threshold hypotheses set includes diurnal, nocturnal, crepuscular, and cathemeral. This set does not define the entire parameter space. The hypotheses of diurnal, nocturnal, and crepuscular are similar to the traditional and general hypotheses sets (using parameter $\xi_1$). The difference is how we define cathemerality, which is done based on the lower threshold value of $\xi_C$. Note that in the `Diel.Niche` package the threshold values are defined for each hypothesis (diurnal, nocturnal, crepuscular, cathemeral) separately as, 'xi.t.D', 'xi.t.N', 'xi.t.CR', 'xi.t.C', respectively.

Researchers may want to use this hypothesis set if they want to consider a traditional hypothesis set but ignore bimodal activity patterns, making the definition of cathemerality more about using all three diel periods a reasonable amount. This hypothesis set is flexible and easily adjusted to change the ranges of each diel phenotype.

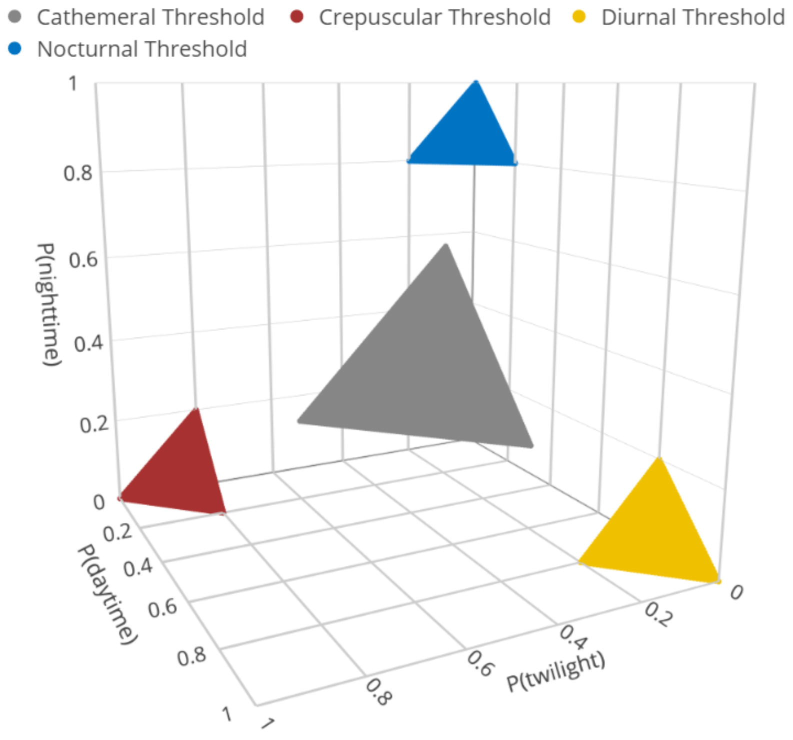


Figure 5: Defined parameter space for the four threshold hypotheses; .

#### 5.1 Diurnal

*Specific Hypothesis*: Probability of activity in the daytime is greater than the probability of activity during the crepuscular and night times. The probability of daytime activity is greater than some threshold ($\xi_D$). There are no constraints on whether probability of nocturnal activity is less than, equal to, or more than crepuscular activity.

*Description*: The probability of twilight and nocturnal activity is lower than the probability of daytime activity, and the probability of daytime activity is greater than $\xi_{\text{D}}$.

*Mathematical Inequalities*:

$$\begin{aligned} p_{\text{tw}} &\leq p_{\text{d}} \\ p_{\text{n}} &\leq p_{\text{d}} \\ \xi_{\text{D}} &\leq p_{d} \end{aligned} \tag{34}$$

*Implementation*: $\boldsymbol{A} = \begin{bmatrix} 1 & -1 \\ -1 & -2 \\ 0 & -1 \end{bmatrix}, \boldsymbol{b} = \begin{bmatrix} 0 \\ -1 \\ -\xi_1 \end{bmatrix}$

Therefore, all together we have,

$$\begin{gathered} \boldsymbol{A\theta} \leq \boldsymbol{b} \\ \begin{bmatrix} (1) \times p_{\text{tw}} + (-1) \times p_{\text{d}} \\ (-1) \times p_{\text{tw}} + (-2) \times p_{\text{d}} \\ (0) \times p_{\text{tw}} + (-1) \times p_{\text{d}} \end{bmatrix} \leq \begin{bmatrix} 0 \\ -1 \\ -\xi_{\text{D}} \end{bmatrix} \\ \begin{bmatrix} p_{\text{tw}} - p_{\text{d}} \\ -p_{\text{tw}} - 2p_{\text{d}} \\ -p_{\text{d}} \end{bmatrix} \leq \begin{bmatrix} 0 \\ -1 \\ -\xi_{\text{D}} \end{bmatrix} \end{gathered} \tag{35}$$

Interpretation Constraint 3:

$$\begin{aligned} &\xi_1 \leq p_{\text{d}} \\ &-p_{\text{d}} \leq -\xi_1, \text{ such that} \\ &(0) \times p_{\text{tw}} + (-1) \times p_{\text{d}} \leq -\xi_{\text{D}} \end{aligned} \tag{36}$$

If $\xi_{\text{D}}$ is $> 0.5$, it will imply that $p_{\text{tw}} < p_{\text{d}}$ and $p_{\text{n}} < p_{\text{d}}$, thus making additional constraints unnecessary.

Interpretation of constraints are the same as the first and second constraints of Maximizing.

#### 5.2 Nocturnal

*Specific Hypothesis*: Probability of activity in the nighttime is greater than the probability of activity during the daytime and the crepuscular times. The probability of nighttime activity is greater than some threshold ($\xi_{\text{N}}$). There are no constraints on whether probability of daytime activity is less than, equal to, or more than crepuscular activity.

*Description*: The probability of twilight and daytime activity is lower than the probability of nighttime activity, and the probability of nighttime activity is greater than $\xi_{\text{N}}$.

*Mathematical Inequalities*:

$$\begin{aligned} p_{\text{d}} &\leq p_{\text{n}} \\ p_{\text{tw}} &\leq p_{\text{n}} \\ \xi_{\text{N}} &\leq p_n \end{aligned} \tag{37}$$

*Implementation*: $\boldsymbol{A} = \begin{bmatrix} 1 & 2 \\ 2 & 1 \\ 1 & 1 \end{bmatrix}, \boldsymbol{b} = \begin{bmatrix} 1 \\ 1 \\ -\xi_{\text{N}} + 1 \end{bmatrix}$

If $\xi_{\text{N}}$ is $> 0.5$, it will imply that $p_{\text{tw}} < p_{\text{n}}$ and $p_{\text{d}} < p_{\text{n}}$, thus making additional constraints unnecessary.

#### 5.3 Crepuscular

*Specific Hypothesis*: The probability of activity in the crepuscular period is greater than the probability of activity during the daytime or nighttime. There is no hypothesized order between activity during the nighttime and daytime. Also, the primary activity during the crepuscular period is greater than the threshold value of $\xi_{\text{CR}}$.

*Description*: The probability of daytime and nighttime activity is lower than the probability of twilight activity, and the probability of twilight activity is greater than $\xi_{\text{CR}}$.

*Mathematical Inequalities*:

$$\begin{aligned} p_{\text{d}} &\leq p_{\text{tw}} \\ p_{\text{n}} &\leq p_{\text{tw}} \\ p_{\text{tw}} &\geq \xi_{\text{CR}} \end{aligned} \tag{38}$$

*Implementation*: $\boldsymbol{A} = \begin{bmatrix} -1 & 1 \\ -2 & -1 \\ -1 & 0 \end{bmatrix}, \boldsymbol{b} = \begin{bmatrix} 0 \\ -1 \\ -\xi_{\text{CR}} \end{bmatrix}$

#### 5.4 Cathemeral

*Specific Hypothesis*: There is a reasonable amount of activity in the daytime, nighttime, and crepuscular periods, such that they are all greater than a certain threshold amount ($\xi_{\text{C}}$).

*Description*: The probability of daytime, nighttime, and twilight activity is all greater than $\xi_{\text{C}}$.

*Mathematical Inequalities*:

$$\begin{aligned} p_{\text{n}} &\geq \xi_C \\ p_{\text{d}} &\geq \xi_C \\ p_{\text{tw}} &\geq \xi_C \end{aligned} \tag{39}$$

$$Implementation: \boldsymbol{A} = \begin{bmatrix} 1 & 1 \\ 0 & -1 \\ -1 & 0 \end{bmatrix}, \boldsymbol{b} = \begin{bmatrix} -\xi_{\mathrm{C}} + 1 \\ -\xi_{\mathrm{C}} \\ -\xi_{\mathrm{C}} \end{bmatrix}$$

### 6 Variation Hypotheses

The variation hypothesis set includes diurnal, nocturnal, crepuscular, and cathemeral. This set does not define the entire parameter space. Rather, we define each diel hypotheses by the lower and upper variation ($\epsilon$) around each most probable value for diurnal, nocturnal, twilight, and cathemeral ($\eta$). Note that in the `Diel.Niche` package the most probable values are defined for each hypothesis (diurnal, nocturnal, crepuscular, cathemeral) separately as, 'eta.D', 'eta.N', 'eta.CR', 'eta.C', respectively.

Researchers may want to use this hypothesis set if they want to consider a traditional hypothesis set but ignores bimodal activity patterns, making the definition of cathemerality more strict as use in all three diel periods is a substantial amount. This setup is similar to the Threshold hypotheses, but is even more flexible in changing the range of probabilities that one can implement for each hypothesis. The Cathemeral Var hypothesis is hexagonal in this set and triangular in the Threshold hypothesis set.

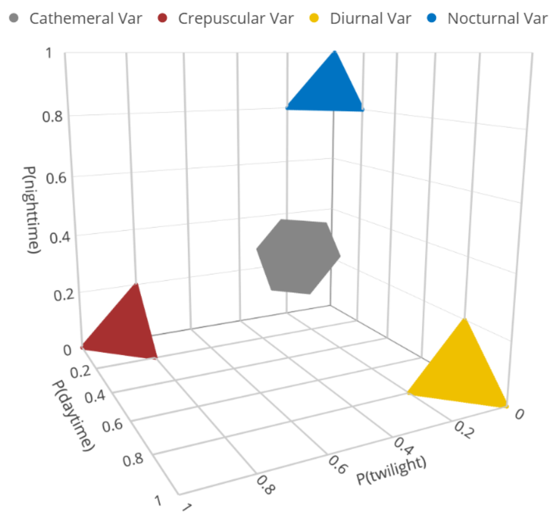


Figure 6: Defined parameter space for the four variation hypotheses; $\eta_{\mathrm{D}}$, $\eta_{\mathrm{CR}}$, $\eta_{\mathrm{N}}$ = 0.9, $\eta_{\mathrm{C}} = 0.33$, and $\epsilon = 0.1$.

#### 6.1 Diurnal

*Specific Hypothesis*: The probability of activity in the daytime is greater than the probability of activity during twilight and nighttime. Also, the probability of daytime activity is in the range of a lower bound of $\eta_{\text{D}} - \epsilon$ and an upper bound of $\eta_{\text{D}} + \epsilon$. There are no constraints on the order between the probability of activity during the nighttime and crepuscular time periods.

*Description*: The probability of twilight and nocturnal activity is lower than the probability of daytime activity, and the probability of daytime activity is in between a lower bound of $\eta_{\text{D}} - \epsilon$ and an upper bound of $\eta_{\text{D}} + \epsilon$.

*Mathematical Inequalities*:

$$\begin{aligned} &p_{\text{tw}} \leq p_{\text{d}} \\ &p_{\text{n}} \leq p_{\text{d}} \\ &\eta_{\text{D}} - \epsilon \leq p_{\text{d}} \leq \eta_{\text{D}} + \epsilon \end{aligned} \tag{40}$$

*Implementation*: $\boldsymbol{A} = \begin{bmatrix} 1 & -1 \\ -1 & -2 \\ 0 & -1 \\ 0 & 1 \end{bmatrix}, \boldsymbol{b} = \begin{bmatrix} 0 \\ -1 \\ -\eta_{\text{D}} + \epsilon \\ \eta_{\text{D}} + \epsilon \end{bmatrix}$

Interpretation of constraints 1 and 2 is the same as for the first and second constraints of Maximizing, while the third constraint can be split into two separate components.

Interpretation of Constraint 3:

$$\begin{aligned} &\eta_{\text{D}} - \epsilon \leq p_{\text{d}} \\ &-p_{\text{d}} \leq -\eta_{\text{D}} + \epsilon, \text{ such that} \\ &(0) \times p_{\text{tw}} + (-1) \times p_{\text{d}} \leq -\eta_{\text{D}} + \epsilon \end{aligned} \tag{41}$$

Interpretation of Constraint 4:

$$\begin{aligned} &p_{\text{d}} \leq \eta_{\text{D}} + \epsilon \\ &(0) \times p_{\text{tw}} + (1) \times p_{\text{d}} \leq \eta_{\text{D}} + \epsilon \end{aligned} \tag{42}$$

#### 6.2 Nocturnal

*Specific Hypothesis*: The probability of activity in the nighttime is greater than the probability of activity during the daytime and crepuscular times. There are no constraints on whether probability of daytime activity is less than, equal to, or more than crepuscular activity. Also, the probability of nighttime activity is in the range of between a lower bound of $\eta_{\text{N}} - \epsilon$ and an upper bound of $\eta_{\text{N}} + \epsilon$.

*Description*: The probability of twilight and daytime activity is lower than the probability of nighttime activity, and the probability of nighttime activity is in between a lower bound of $\eta_\mathrm{N} - \epsilon$ and upper bound of $\eta_\mathrm{N} + \epsilon$.

*Mathematical Inequalities*:

$$\begin{aligned} & p_\mathrm{d} \leq p_\mathrm{n} \\ & p_\mathrm{tw} \leq p_\mathrm{n} \\ & \eta_\mathrm{N} - \epsilon \leq p_\mathrm{n} \leq \eta_\mathrm{N} + \epsilon \end{aligned} \tag{43}$$

*Implementation*: $\boldsymbol{A} = \begin{bmatrix} 1 & 2 \\ 2 & 1 \\ -1 & -1 \\ 1 & 1 \end{bmatrix}, \boldsymbol{b} = \begin{bmatrix} 1 \\ 1 \\ \eta_\mathrm{N} + \epsilon - 1 \\ -\eta_\mathrm{N} + \epsilon + 1 \end{bmatrix}$

#### 6.3 Crepuscular

*Specific Hypothesis*: The probability of activity in the crepuscular period is greater than the probability of activity during the daytime or nighttime. There is no hypothesized order between activity during the nighttime and daytime. Also though, the probability of crepuscular activity is in the range of between a lower bound of $\xi_3 - \epsilon$ and an upper bound of $\xi_3 + \epsilon$.

*Description*: The probability of nighttime and daytime activity is lower than the probability of twilight activity, and the probability of twilight activity is in between a lower bound of $\eta_\mathrm{CR} - \epsilon$ and upper bound of $\eta_\mathrm{CR} + \epsilon$.

*Mathematical Inequalities*:

$$\begin{aligned} & p_\mathrm{d} \leq p_\mathrm{tw} \\ & p_\mathrm{n} \leq p_\mathrm{tw} \\ & \xi_3 - \epsilon \leq p_\mathrm{tw} \leq \eta_\mathrm{CR} + \epsilon \end{aligned} \tag{44}$$

*Implementation*: $\boldsymbol{A} = \begin{bmatrix} -1 & 1 \\ -2 & -1 \\ -1 & 0 \\ 1 & 0 \end{bmatrix}, \boldsymbol{b} = \begin{bmatrix} 0 \\ -1 \\ -\eta_\mathrm{CR} + \epsilon \\ \eta_\mathrm{CR} + \epsilon \end{bmatrix}$

#### 6.4 Cathemeral

*Specific Hypothesis*: Daytime, nighttime, and crepuscular activity is similar and in the range between a lower bound of $\eta_\mathrm{C} - \epsilon$ and an upper bound of $\eta_\mathrm{C} + \epsilon$.

*Description*: The probability of nighttime, daytime, and twilight activity is in between a lower bound of $\eta_\mathrm{C} - \epsilon$ and upper bound of $\eta_\mathrm{C} + \epsilon$.

*Mathematical Inequalities*:

$$\begin{aligned} \xi_C - \epsilon \leq p_{\mathrm{tw}} \leq \eta_{\mathrm{C}} + \epsilon \\ \xi_C - \epsilon \leq p_{\mathrm{d}} \leq \eta_{\mathrm{C}} + \epsilon \\ \xi_C - \epsilon \leq p_{\mathrm{n}} \leq \eta_{\mathrm{C}} + \epsilon \end{aligned} \tag{45}$$

*Implementation*: $\boldsymbol{A} = \begin{bmatrix} 1 & 0 \\ -1 & 0 \\ 0 & 1 \\ 0 & -1 \\ -1 & -1 \\ 1 & 1 \end{bmatrix}, \boldsymbol{b} = \begin{bmatrix} \eta_{\mathrm{C}} + \epsilon \\ -\eta_{\mathrm{C}} + \epsilon \\ \eta_{\mathrm{C}} + \epsilon \\ -\eta_{\mathrm{C}} + \epsilon \\ \eta_{\mathrm{C}} + \epsilon - 1 \\ -\eta_{\mathrm{C}} + \epsilon + 1 \end{bmatrix}$

### 7 Additional Hypotheses

We've included a few extra hypotheses here that may be useful in special cases. They can be included in combination with other hypotheses from the sets defined above.

#### 7.1 Unconstrained Hypothesis

*Specific Hypothesis*: There are no constraints on an animal's activity. Researchers may want to consider this hypothesis to evaluate whether constraints improve parameter estimation. Or, when using a hypothesis set that has certain probability spaces excluded (Variation and Threshold), this hypothesis could be used as a catch all for the spaces that are not of interest.

*Description*: The probability of activity during twilight, nighttime, and daytime are between 0 and 1.

*Mathematical Inequalities*:

$$\begin{aligned} p_{\mathrm{d}} \leq 1 \\ p_{\mathrm{d}} \geq 0 \\ p_{\mathrm{tw}} \leq 1 \\ p_{\mathrm{tw}} \geq 0 \end{aligned} \tag{46}$$

*Implementation*: $\boldsymbol{A} = \begin{bmatrix} 0 & 1 \\ 0 & -1 \\ 1 & 0 \\ -1 & 0 \end{bmatrix}, \boldsymbol{b} = \begin{bmatrix} 1 \\ 0 \\ 1 \\ 0 \end{bmatrix}$

#### 7.2 Available Cathemeral Variation Hypothesis

*General Hypothesis*: An animal is cathemeral and active during all time periods in similar proportion to amount of available time in each period.

This hypothesis is specifying that diel periods are used in proportion to their availability, but allows some variation or error around this. It is the same idea as the Selection hypothesis specified in section 4.7. However, that hypothesis is specified using an equality constraint and is thus much more precise. This makes the sample size requirements much larger to identify this model as supported in a set. However, this hypothesis relaxes the constraint, which makes it more realistically usable.
We require proportional availability by time category to be supplied to evaluate this hypothesis. Let's define $\mathbf{p}_{\text{avail}} = [p_{\text{av.tw}},\ p_{\text{av.d}}]$. For example, $\mathbf{p}_{\text{avail}} = [0.16\ \ 0.44]$.
*Specific Hypothesis*: The probability of activity in each time period divided by the available time is within the range of $1 - \epsilon \leq p \leq 1 + \epsilon$.

*Mathematical Inequalities*:

$$\begin{aligned} 1 - \epsilon &\leq p_{\text{tw}}/p_{\text{av.tw}} \leq 1 + \epsilon \\ 1 - \epsilon &\leq p_{\text{d}}/p_{\text{av.d}} \leq 1 + \epsilon \\ 1 - \epsilon &\leq p_{\text{n}}/p_{\text{av.n}} \leq 1 + \epsilon \end{aligned} \tag{47}$$

*Implementation*: $\boldsymbol{A} = \begin{bmatrix} -1/p_{\text{av.tw}} & 0 \\ 1/p_{\text{av.tw}} & 0 \\ 0 & -1/p_{\text{av.d}} \\ 0 & 1/p_{\text{av.d}} \\ 1/(1 - p_{\text{av.tw}} - p_{\text{av.d}}) & 1/(1 - p_{\text{av.tw}} - p_{\text{av.d}}) \\ -1/(1 - p_{\text{av.tw}} - p_{\text{av.d}}) & -1/(1 - p_{\text{av.tw}} - p_{\text{av.d}}) \end{bmatrix}, \boldsymbol{b} = \begin{bmatrix} -1 + \epsilon \\ 1 + \epsilon \\ -1 + \epsilon \\ 1 + \epsilon \\ -1 + \epsilon + 1/p_{\text{av.n}} \\ 1 + \epsilon - 1/p_{\text{av.n}} \end{bmatrix}$

#### 7.3 Even Cathemeral Hypothesis

*General Hypothesis*: An animal is cathemeral and active evenly throughout the 24-hour period.

A strict definition of cathemerality is that all three diel periods are equal, such that an even amount of activity occurs throughout the entire 24-hr period. This hypothesis specifies this idea. However, because it's an equality there is very little parameter space and thus a large sample size is needed.

##### 7.3.1 Equal

*Specific Hypothesis*: Daytime, nighttime, and crepuscular activity are equal with a probability of 0.3333.

*Mathematical Inequalities*:

$$p_{\text{tw}} = p_{\text{d}} = p_{\text{n}} = 0.33 \tag{48}$$

Implementation: $\boldsymbol{C} = \begin{bmatrix} 1 & 0 \\ 0 & 1 \end{bmatrix}, \boldsymbol{d} = \begin{bmatrix} 0.33 \\ 0.33 \end{bmatrix}$

### References

Heck, D. W., & Davis-Stober, C. P. (2019). Multinomial models with linear inequality constraints: Overview and improvements of computational methods for Bayesian inference. Journal of mathematical psychology, 91, 70-87.
